# Kv1.3 palmitoylation regulates spatial distribution and channel removal from the immunological synapse

**DOI:** 10.64898/2026.01.19.700329

**Authors:** María Navarro-Pérez, Mireia Pérez-Verdaguer, Anna Benavente-Garcia, Michael L Dustin, Antonio Felipe, Jesusa Capera

**Affiliations:** Departament de Bioquímica i Biomedicina Molecular, Institut de Biomedicina (IBUB), Universitat de Barcelona, Barcelona, Spain; The Kennedy Institute of Rheumatology, Nuffield Department of Orthopaedics Rheumatology & Musculoskeletal Sciences, University of Oxford, Oxford, United Kingdom; Department of Biology, University of Padova, Padova, Italy

**Keywords:** Potassium channels, Palmitoylation, Ubiquitination, Immunological synapse, Lymphocytes

## Abstract

Kv1.3 is the main voltage-gated potassium channel in T cells. At the immunological synapse (IS), it sustains Ca²⁺ signaling and facilitates T cell activation. Aberrant Kv1.3 expression or activity is linked to autoimmune disorders, yet the mechanisms regulating its targeting and organization at the IS remain unclear. We show that Kv1.3 palmitoylation is a dynamic process mediating channel rearrangement at the IS. The ZDHHC21 acyltransferase, which also S-acylates the TCR, palmitoylates Kv1.3, positioning this enzyme as a potential therapeutic target. Palmitoylation promotes channel migration to the synapse center for removal from the surface. A nonpalmitoylated mutant (Cys*_less_* Kv1.3) accumulated at the distal IS and was excluded from lipid raft–enriched domains. Mislocalization and reduced current hindered lymphocyte activation. Moreover, Cys*_less_* Kv1.3 showed stronger interaction with PSD95 and cortactin, stabilizing the channel at the surface. These findings highlight S-palmitoylation as a crucial regulator of Kv1.3 during immune responses and a promising target in autoimmune disease therapy.

## Introduction

Leukocyte physiology relies on the activity of a limited repertoire of ion channels that, by initiating Ca^2+^ signaling, govern the immune response. In this context, the voltage-gated potassium channel Kv1.3 is essential for generating K^+^ efflux that facilitates sustained Ca^2+^ entry through CRAC channels upon TCR engagement [1]. This signaling process is concentrated mainly at specific locations at the interface between T cells and antigen-cells (APCs), where antigen presentation triggers the formation of the immunological synapse (IS) [2].

The IS is a highly organized structure comprising concentric supramolecular activation clusters (SMACs). The central SMAC (cSMAC) segregates the TCR-CD3 complex and coreceptors CD4 or CD8 upon interaction with the major histocompatibility complex (MHC) on an APC and serves as a site of active internalization and release of synaptic components in extracellular vesicles. Additionally, CD28, a costimulatory receptor, enhances TCR signaling by interacting with CD80 or CD86 on the APC. Surrounding the cSMAC is the peripheral SMAC (pSMAC), which contains adhesion molecules such as lymphocyte function-associated antigen-1 (LFA1). LFA-1 interacts with intercellular adhesion molecule-1 (ICAM-1) on APCs, strengthening cell interactions. The outermost region, named the distal SMAC (dSMAC), supports dynamic processes such as synapse projection and retraction. dSMAC is enriched in CD45, a receptor-like protein tyrosine phosphatase that is critical for regulating signal transduction pathways. Other receptors, such as CD2 and CD58, also contribute to the structural and functional integrity of the dSMAC [2,3].

Kv1.3 is primarily localized at the dSMAC, and colocalizes with the CD2/CD58 corolla, which increases TCR-dependent Ca^2+^ signaling [4]. Moreover, the IS is enriched in lipid raft microdomains, where Kv1.3 is partially localized [5–9]. Upon synapse maturation, the channel steadily migrates to the cSMAC, where it undergoes endocytosis [4]. While Kv1.3 reaches the IS by lateral diffusion at the cell surface [10], channel trafficking to the plasma membrane mostly relies on a diacidic COPII signal at the C-terminal domain [11]. Annihilation of this domain dramatically impairs Kv1.3 surface expression, recruitment into the IS, and the association of membrane-associated guanylate kinases (MAGUK) [11–13]. MAGUK proteins, such as PSD-95, facilitate and stabilize Kv1.3 polarization into the IS [12,13]. Kv1.3 targets lipid rafts via its association with caveolins. However, T lymphocytes largely lack caveolin, leaving the mechanisms of Kv1.3 recruitment to LR uncertain [14]. In this context, many membrane proteins target the plasma membrane via lipidic posttranslational modifications such as S-acylation. For instance, the regulatory Kvβ2.1 subunit, despite being a cytosolic protein, reaches lipid raft microdomains and the IS undergoing palmitoylation. This process occurs independently of caveolin association and facilitates membrane targeting during lymphocyte activation [15]. Palmitoyl acyltransferases catalyze the binding of palmitate to cysteine residues, which can be reverted by thioesterases. The reversible nature of this posttranslational modification positions it as a key regulator of TCR-mediated signaling during lymphocyte activation. TCR stimulation triggers the palmitoylation of numerous proteins involved in cell signaling, vesicle transport, and adaptor functions [16,17]. Consequently, as palmitoylation has emerged as a pivotal modulator of immune sensing and signaling, recent studies have linked its dysregulation to various immunological disorders, highlighting its potential as a therapeutic target [18]. Therefore, palmitoylation, which facilitates protein–protein interactions, increases protein stability, and improves trafficking and membrane association [19,20], is worthy of investigation, as it has emerged as a key regulatory mechanism of leukocyte physiology.

Our study demonstrates, for the first time, that the palmitoylation of Kv1.3 regulates the spatial distribution of the channel within SMAC compartments at the IS. Kv1.3 is palmitoylated by ZDHHC21 acyltransferase at multiple intracellular cysteines. Palmitoylation promotes Kv1.3 endocytosis at the cSMAC, thus regulating channel abundance at the synapse and termination of its function. Furthermore, Kv1.3 palmitoylation promotes T-cell activation by promoting early TCR signaling and Ca^2+^ flux at the IS. Our findings reveal that palmitoylation is a dynamic posttranslational modification that regulates membrane arrangement and function at the IS platform. Moreover, our work expands the repertoire of strategies to target Kv1.3-related autoimmune disorders by introducing the functional implications of channel palmitoylation in the complex scenario of the immune system response.

## Materials and methods

### Expression plasmids and site-directed mutagenesis

T.C. Holmes (University of California, Irvine, CA) provided the rat Kv1.3 in the pRcCMV construct. The human Kv1.3 pBSTA plasmid was kindly provided by F. Bezanilla (University of Chicago, IL). The channels were subcloned and inserted into pEYFP-C1 plasmids (Clontech). All Kv1.3 mutants were generated using a QuikChange multisite-directed mutagenesis kit (Agilent Technologies). For mRNA electroporation, Kv1.3 pEYFP-C1 and its mutants were subcloned and inserted into pcDNA3 with a T7 promoter. All the constructs were verified by sequencing.

### Cell culture, isolation of T-cell subsets and blast generation

HEK293 cells were cultured in Dulbecco’s modified Eagle’s medium (DMEM; Gibco) supplemented with 4.5 g/L glucose and L-glutamine and supplemented with 10% FBS (fetal bovine serum; Gibco) and antibiotics (10000 U/mL penicillin G, 10 mg/mL streptomycin; Gibco). Lipotransfectin® (AttendBio Research) was used for DNA transfection when the cells were approximately 80% confluent. The cells were subsequently starved overnight (O/N) with DMEM supplemented with 0.2% bovine serum albumin (BSA; Sigma‒Aldrich).

Human T-cell subsets were isolated from anonymized blood samples provided by the National Health Service Blood and Transplant (UK) and Banc de Sang I Teixits de Catalunya (Spain) using negative selection kits from Stemcell Technologies. Rosette Sep (Stemcell Technologies) was used to enrich total CD4^+^ T cells. Human T cells were cultured at 37 °C and 5% CO_2_ in RPMI 1640 medium (Life Technologies) supplemented with 10% FBS, 1% glutamine, 1% penicillin‒streptomycin (Gibco), 1x nonessential amino acid solution (Thermo Fisher Scientific), 10 mM HEPES (Life Technologies) and 50 U/ml IL-2. To generate T-cell blasts, the Dynabeads™ Human T-Activator CD3/CD28 for T-cell expansion and activation kit (Life Technologies) was used following the manufacturer’s instructions. Human T-cell blasts were ready for use in experiments after 6–7 days of expansion. No IL-2 was supplemented in the media the day before the experiment. For the culture of the Jurkat human T-cell line and the Raji human B-cell line, RPMI 1640 medium supplemented with 10% FBS and 1% penicillin‒streptomycin was used.

### RNA transfection of human T cells

RNA transcripts were prepared from DNA constructs in pcDNA3 plasmids linearized with NdeI restriction enzyme at 37 °C for 2 h. DNA precipitation, in vitro RNA transcription and in vitro polyadenylation were performed following the protocol of the mMESSAGE mMACHINE T7 ULTRA transcription kit (Thermo Fisher Scientific). T cells in culture were washed three times in Opti-MEM (Life Technologies) and resuspended at 2–2.5x10^6^ cells per 100 μl. Each 100 μl cell suspension was mixed with the RNA preparation and transferred to a gene pulser/micropulser electroporation cuvette (0.2 cm gap; Bio-Rad). Cells were electroporated at 300 V for 2 ms in an ECM 830 square wave electroporation system (BTX). The cells were incubated at 2x10^6^ cells/ml for at least 18 h at 37 °C in a 5% CO_2_ incubator before the experiments were performed.

### siRNA transfection

First, 3.3 nmol of lyophilized ZDHHC21 siRNA (Santa Cruz Biotechnology) was reconstituted in 330 μL of RNAse-free water to a final concentration of 10 μM. Next, human primary CD4^+^ T cells were electroporated with 10 μL of siRNA stock as indicated above. Simultaneously, 10 μL of the scrambled sequence control siRNA-A (Santa Cruz Biotechnology) was used as a mock siRNA for the negative control. Cells were incubated in complete media at 37 °C and 5% CO_2_ for 24 h. Then, gene silencing was verified by protein extraction and Western blot analysis. After positive results were obtained, the experiment was conducted.

### Supported lipid bilayer (SLB), immunostaining, and TIRFM imaging

SLB were prepared as previously described [21]. Briefly, glass coverslips (Nexterion) were cleaned in piranha solution, rinsed in water, dried, plasma-cleaned and mounted onto six-channel chambers (Ibidi). Small unilamellar liposomes were prepared by extrusion using 100 mol% 1,2-dioleoyl-sn-glycero-3-phosphocholine (Avanti Polar Lipids Inc.) or 75% 1,2-dioleoyl-sn-glycero-3-phosphocholine and 25% 1,2-dioleoyl-sn-glycero-3-[(N-(5-amino-1-carboxypentyl) iminodiacetic acid) succinyl]-Ni (Avanti Polar Lipids Inc.) at a final lipid concentration of 0.4 mM. The channels in the Ibidi chamber were covered with a 1:1 mixture of these two liposome preparations, blocked and washed. Channels in the Ibidi chamber were covered with a liposome mixture, and after a 20 min incubation at room temperature (RT), they were washed with phosphate-buffered saline (PBS). SLBs were blocked with 3% bovine serum albumin (BSA)-supplemented PBS supplemented with 100 μM NiSO_4_ to saturate the NTA sites for 20 min. Combinations of the following reagents were then incubated onto the SLBs at specific concentrations to achieve the indicated molecular densities: His-tagged UCHT1 (30 molecules/μm^2^), ICAM1 (200 molecules/μm^2^), CD58 (200 molecules/μm^2^), and CD80 (200 molecules/μm^2^). The channels were subsequently washed and kept in PBS until use. Cells were added to the SLBs and immediately imaged. For lipid raft staining, lymphocytes were previously incubated with cholera toxin subunit B (Fisher Scientific). For the fixed experiments, the cells were allowed to interact with SLBs for 15 min at 37 °C before they were fixed for 10 min with 4% PFA in PHEM buffer at 37 °C. The cells were blocked in 3% BSA supplemented with 100 mM Gly for 30 min at RT. Cells were stained for 30 min at RT with an Alexa Fluor® 647-conjugated anti-ZAP-70 antibody (BioLegend) when indicated. Imaging was performed on an Olympus IX83 inverted microscope equipped with a TIRF module. The instrument was equipped with an Olympus UApON 150x TIRF N.A. 1.45 objective; 405, 488, 568, and 640 nm laser lines; and a Photometrics Evolve delta EMCCD camera. In live imaging experiments, SLBs were transferred to a preheated incubator on top of the TIRF microscope, and cells were added to the well. TIRF images were analyzed using Fiji (ImageJ). For single-particle tracking (SPT) analysis, the TrackMate plugin v7.6.1 [22] was used on videos captured at 0.3 sec/frame for 1 min. First, particles were detected with an estimated diameter of 0.3 μm. A linear assignment problem tracker was used for the frame-to-frame particle linking with the following parameters: a maximum linking distance of 0.3 μm, a gap-closing maximum distance of 0.4 μm, and a gap-closing maximum frame gap of 1 frame.

### Immunocytochemistry, cell conjugates, and confocal imaging

HEK293 cells were plated on polylysine-coated coverslips. Twenty-four hours after transfection, the cells were washed with PBS and fixed for 10 min at RT with 4% PFA (paraformaldehyde; Sigma‒Aldrich). For plasma membrane labeling, the cells were incubated for 5 min on ice with Alexa Fluor® 555 WGA (Wheat Germ Agglutinin; Fisher Scientific) prior to fixation. To study endocytosis, the samples were incubated in blocking solution (5% nonfat milk and 0.5% Triton X-100 in PBS) for 1 h at RT, after which they were incubated with anti-EEA1 primary antibodies (Fisher Scientific) for 2 h at RT. Alexa Fluor® 647-conjugated goat anti-mouse secondary antibodies were incubated for 2 h at RT, and the samples were mounted on slides with homemade Mowiol mounting solution.

For the cell conjugates, Raji human B cells were used as antigen-presenting cells (APC). Raji cells were prepulsed with 1 μg/ml of the superantigen Staphylococcal enterotoxin B (SEB) for 30 min at 37 °C in a 5% CO_2_ incubator. After two washes with RPMI 1640 media, the prepulsed Raji B cells were mixed with human CD4^+^ T cells at a 1:1 ratio for 15 min at 37 °C in a 5% CO_2_ incubator. The mixtures were fixed with 2% PFA for 10 min at RT. Conjugates were plated on poly-L-lysine-coated coverslips and blocked with 3% BSA supplemented with 100 mM Gly for 1 h at RT. Next, the cells were labeled with primary anti-CD3 (Alexa Fluor 647; Biolegend), anti-CD19 (Brilliant Violet 421; Biolegend), anti-ZDHHC21 (NSJ Bioreagents) and anti-Kv1.3 (FITC; Alomone) antibodies in 1% BSA for 1 h at RT. After three washes with PBS, the coverslips were mounted in Mowiol (Calbiochem). Confocal images were acquired with a Zeiss 880 confocal microscope. The images were analyzed using Fiji (ImageJ).

### Click-chemistry and proximity ligation assay (PLA)

Primary human CD4^+^ T cells were incubated with 100 μM sonicated 15-hexadecynoic palmitic acid (15-yne, Avanti Polar Lipids) for 18 h at 37 °C, enabling Alk-C16 protein palmitoylation. Next, the cells were washed, and immunological synapses were generated with either Raji B cells or SLB. After 15 min of incubation at 37°C, the lymphocytes were fixed with 2% PFA followed by cold methanol for 5 min. The cells were rinsed in PBS, and the click reaction (0.1 mM biotin-azide (carboxamide-6-azidohexanyl biotin, Invitrogen), 0.1 mM TCEP (Tris(2-carboxyethyl)phosphine hydrochloride, Sigma‒Aldrich) and 0.1 mM CuSO_4_ (Sigma‒Aldrich)) was incubated with gentle rocking for 1 h at RT. After five washes with PBS without K^+^, the cells were incubated with blocking solution (5% BSA and 0.3% Triton X-100) for 1 h at RT. The lymphocytes were further washed three times with K^+^-free PBS and incubated with primary antibodies diluted in 1X Duolink blocking solution (Sigma‒Aldrich) at 4 °C overnight. Primary antibodies included either monoclonal anti-Kv1.3 (1/50, Neuromab) for native Kv1.3 or monoclonal anti-GFP (1/100, Roche) for Kv1.3-electroporated CD4^+^ T cells. For palmitoylation detection, polyclonal antibiotin (1/300, Sigma‒Aldrich) was used to detect the biotin linked to the palmitic acid, whereas for ubiquitination detection, polyclonal anti-ubiquitin (1/100, CliniSciences) was used. After 3 quick washes, the secondary PLA antibodies were incubated for 1 h at 37 °C (100 μL: 20 μL of Duolink in situ PLA probe anti-Mouse PLUS, 20 μL of Duolink in situ PLA probe anti-Goat MINUS and 60 μL of 1x diluent Duolink antibodies; Sigma‒Aldrich). The cells were subsequently rinsed 5 times and incubated for 30 min at 37 °C with PLA ligation solution (100 μL of 20 μL of 5x ligation buffer, 2.5 μL of ligase and 77.5 μL of Milli-Q® water; Duolink Ligase; Sigma‒Aldrich). After 5 washes, the lymphocytes were incubated with the PLA amplification solution (100 μL: 20 μL of 5x amplification buffer, 1.25 μL of polymerase and 78.75 μL of Milli-Q® water; Duolink Far Red amplification, Sigma‒Aldrich) at 37 °C for 3 h in the dark. Next, the cells were further washed 4 times and incubated with anti-Kv1.3 conjugated with Fitc (1/100, Alomone) for 2 h at RT to label total Kv1.3. Finally, the samples were observed under an inverted Zeiss LSM880 laser scanning confocal microscope.

### Ca^2+^ imaging

Human primary CD4⁺ T cells were loaded with 5 μM Calbryte™ 630 AM in Hank’s balanced salt solution with 20 mM HEPES (HHBS) supplemented with 0.04% Pluronic for 1 h at 37 °C. Following two washes with HHBS, fluorescence was recorded using the Zeiss LSM880 confocal microscope. Videos of immunological synapse formation with Raji B cells were recorded at 0.1 s/frame for 1 min. Image analysis was conducted using ImageJ software.

### Flow cytometry

Human primary CD4^+^ T cells were electroporated with either Kv1.3-YFP wild-type (WT) or Cys mutant constructs and incubated O/N at a 1:1 ratio with anti-CD3/CD28 Dynabeads (Gibco) at 37 °C. Following incubation, the cells were washed and fixed with 2% PFA for 15 min at RT. Fixed cells were then washed three times with PBS and incubated with Pacific Blue™ anti-human CD279 (PD-1) antibody (BioLegend), APC anti-human CD69 antibody (BioLegend), and PE anti-human Cd25 (BioLegend) for 2 h at 4 °C. The cells were subsequently washed three times and analyzed using a Gallios flow cytometer (Beckman Coulter). Analysis was restricted to YFP-positive cells. The data were processed and analyzed using FlowJo software (version 10.7.1).

### Protein extraction, plasma membrane purification, immunoprecipitation, ubiquitination, and membrane protein biotinylation

HEK293 cells were washed twice with cold PBS and collected by scraping in 0.5 mL of protein lysis buffer (150 mM NaCl, 50 mM Tris-HCl, 1 mM EDTA, and 1% Triton X-100, pH 7.5) supplemented with protease inhibitors (1 μg/mL aprotinin, 2 μM leupeptin, 1 μM pepstatin, and 1 mM PMSF). The lysates were homogenized by incubation on an orbital shaker at 4 °C for 10 min, followed by centrifugation at 15,000 × g for 15 min. The supernatants were collected, whereas the pellets containing cell debris and nonlysed cells were discarded. The protein concentration was determined using the Bradford protein assay.

Plasma membrane fractions were purified from transfected HEK293 cells using a differential centrifugation protocol adapted from [23]. Briefly, cells were trypsinized, washed twice with Ca^2+^-free PBS, and centrifuged at 600 × g for 10 min. The cell pellet was homogenized in 2 mL of Buffer 1 (225 mM mannitol, 75 mM sucrose, 0.1 mM EGTA, and 30 mM Tris, pH 7.4) and centrifuged at 600 × g for 10 min to remove cellular debris and nuclei. The resulting supernatant was centrifuged at 7000 × g for 10 min to eliminate additional debris. The supernatant from this step was further centrifuged at 20,000 × g for 30 min to isolate a membrane-enriched pellet. The membranous fractions were resuspended in 50 μL of Buffer 2 (225 mM mannitol, 75 mM sucrose, and 30 mM Tris, pH 7.4). For Western blot analysis, all samples were mixed with Laemmli buffer containing 2% β-mercaptoethanol and boiled.

Coimmunoprecipitation experiments were conducted using HEK293 cells 24 h posttransfection. The cells were subsequently washed twice with ice-cold PBS, after which the protein was extracted as previously described. Protein A-Sepharose beads (GE Healthcare) were incubated with an anti-Kv1.3 antibody (Alomone) for 1 h at RT to form bead‒antibody complexes. These complexes were covalently cross-linked using 20 mM dimethyl pimelimidate (DMP) (Pierce) for 30 min at RT. The reaction was quenched with 0.2 M glycine (pH 2.5). To reduce nonspecific binding, cell lysates were precleared by incubation with Protein A-Sepharose beads for 1 h at 4 °C with gentle mixing. Equal amounts of protein (1–2 mg) were then incubated with the bead–antibody complexes O/N at 4 °C. After five washes, the bound proteins were eluted in 100 μL of 0.2 M glycine (pH 2.5).

To study Kv1.3 ubiquitination, the cells were washed twice with ice-cold PBS, and the dishes were frozen O/N. The cells were then lysed for 20 min in lysis buffer supplemented with 10 nM N-ethylmaleimide (NEM), 0.2 mM MG-132, 1 mM EGTA, 1 mM EDTA, 20 mM NaF, 1 mM Na₃VO₄, 2 mM DTT, and 1% Triton X-100.

For immunoprecipitation, 1–2 mg of total protein was precleared by incubation with 30 μL of Protein A-Sepharose™ 4 Fast Flow beads (GE Healthcare) for 1 h at 4 °C with gentle mixing. The beads were removed by centrifugation at 1000 × g for 30 s. The precleared samples were then incubated O/N at 4 °C in Micro Bio-Spin™ Chromatography Columns (Bio-Rad) with 50 μL of Protein A-Sepharose beads and anti-GFP antibody (4 ng antibody/μg protein; GenScript) with gentle agitation. After centrifugation at 1000 × g for 30 s, the supernatant was collected, and the beads were washed five times with washing buffer (150 mM NaCl, 50 mM HEPES, 10% glycerol, 0.1% Triton X-100; pH 7.5). Immunoprecipitates were eluted in 60 μL of 0.2 M glycine (pH 2.5). Laemmli loading buffer containing β-mercaptoethanol was added, and the samples were boiled for 7 min before further analysis.

### Raft isolation and Western blotting

Low-density, Triton-insoluble complexes were isolated as previously described. Briefly, after three washes in PBS, the cells were homogenized in 1 ml of 1% Triton X-100 MBS (150 mM NaCl, 25 mM 2-morpholinoethanesulfonic acid 1-hydrate (MES), pH 6.5) supplemented with 1 μg/mL aprotinin, 1 μg/ml leupeptin, 1 μg/mL pepstatin and 1 mM phenylmethylsulfonyl fluoride (PMSF) to inhibit proteases. Sucrose in MBS was added to a final concentration of 40%. A 5–30% linear sucrose gradient was layered on top and further centrifuged (39000 rpm) for 20–22 h at 4 °C in a Beckman SW41Ti swinging rotor. Gradient fractions (1 ml) were collected from the top, and 50 μL of each raft fraction was boiled in Laemmli SDS loading buffer and separated by 10% SDS‒PAGE. Next, the samples were transferred to PVDF membranes (Immobilon-P, Millipore) and blocked with 5% dry milk supplemented with 0.05% Tween-20 in PBS. The filters were then immunoblotted with specific antibodies: anti-Kv1.3 (1:200; Neuromab), anti-clathrin (1:1000; BD Biosciences), and anti-flotillin (1:500; BD Biosciences). Finally, the filters were washed with 0.05% Tween-20 in PBS and incubated with horseradish peroxidase-conjugated secondary antibodies (Bio-Rad).

### Acyl-biotinyl exchange (ABE) method

Protein palmitoylation was detected using the ABE method adapted from [24]. Briefly, cells were washed with PBS and lysed in lysis buffer (50 mM Tris, 0.2% Triton X-100, 5 mM EDTA, 150 mM NaCl; pH 7.4) supplemented with 1 mg/ml aprotinin, 1 mg/ml leupeptin, 1 mg/ml pepstatin, and 1 mM PMSF as protease inhibitors. To block free thiols, 10 mM N-ethylmaleimide (NEM; Sigma) was added. The samples were incubated at 4 °C for 1 h. The lysates were subsequently cleared by centrifugation at 16000 × g for 15 min, after which the resulting supernatants were used for the ABE assay. The proteins in the supernatants were precipitated with chloroform–methanol (CM), and the protein pellets were solubilized in 0.3 ml of resuspension buffer (50 mM Tris, 4% SDS, pH 7.4) supplemented with 10 mM NEM. Next, 0.9 ml of lysis buffer supplemented with 1 mM NEM was added. The samples were incubated at 4 °C O/N. Excess NEM was removed by three sequential CM precipitations followed by solubilization in 0.5 ml of resuspension buffer. To cleave thioester bonds and allow incorporation of a biotin moiety at exposed sulfur atoms, each sample was diluted in HA buffer (50 mM Tris, 0.2% Triton X-100, pH 7.4) supplemented with protease inhibitors, 1 mM EZ-link HPDP-biotin (ThermoScientific) and 0.7 M hydroxylamine (NH_2_OH) for HA+ samples. As a negative control, a duplicate sample was incubated in the absence of NH_2_OH (HA-). In this case, palmitate groups are not removed, thus preventing biotinylation-mediated purification. The mixture was incubated at RT for 1 h and subjected to CM precipitation. The precipitated protein was solubilized in 0.3 ml of resuspension buffer, diluted with 0.9 ml of lysis buffer containing 0.2 mM HPDP-biotin and protease inhibitors, and incubated for 1 h at RT. Unreacted HPDP-biotin was removed by three sequential CM precipitations, and the protein pellets were solubilized in 0.1 ml of final buffer (50 mM Tris, 0.2% Triton X-100, 5 mM EDTA, 2% SDS, pH 7.4). The samples were diluted in 2 mL of lysis buffer containing protease inhibitors and incubated at RT for 30 min. The samples were briefly centrifuged, after which the nonsuspended particles were discarded. A 40 μL sample was collected as the starting material (SM). Surplus samples were incubated with 50 μL of precleaned NeutrAvidin Agarose beads (Thermo Scientific) O/N at 4 °C. After the beads were washed with lysis buffer, the bound proteins were eluted with 50 μL of SDS‒PAGE sample buffer at 95 °C for 7 min. The eluates (pull-downs) and SM were subjected to SDS‒PAGE and analyzed by Western blotting. To quantify protein palmitoylation, the pulldown band intensity was normalized to that of SM.

### Electrophysiology

The cells were trypsinized and replated onto a perfusion chamber. After a 15-min incubation, the cells were thoroughly washed with extracellular solution containing (in mM) 145 NaCl, 4 KCl, 1 MgCl₂, 1.8 CaCl₂, 10 HEPES, and 10 glucose, with the pH adjusted to 7.4 using NaOH. Micropipettes were fabricated from borosilicate glass capillaries (1.2 mm outer diameter × 0.944 mm inner diameter × 100 mm length; Harvard Apparatus) using a P-97 puller (Sutter Instruments) and fire-polished with an MF-830 Microforge (Narishige) to achieve a resistance of 2–5 MΩ. The intracellular pipette solution for HEK293 cells contained (in mM) 80 aspartate potassium (AspK), 42 KCl, 10 KH₂PO₄, 5 EGTA-K, 5 HEPES-K (pH 7.0 adjusted with KOH), 3 phosphocreatine, and 3 ATP-Mg. The cells were voltage clamped at a holding potential of -80 mV. Potassium current recordings were performed at RT using the whole-cell configuration of the patch-clamp technique with a HEKA EPC-10 amplifier (Harvard Bioscience, Holliston, MA, USA) and the acquisition software PATCHMASTER v2x91 (HEKA Elektronik GmbH). A 45-s interval between pulses was maintained to allow complete recovery of Kv1.3 from inactivation. Voltage-gated currents were elicited by stimulating cells with 250 ms square pulses at +60 mV.

### Statistics

The results are expressed as the mean ± SE. Student’s *t* test, paired *t* test, one-way ANOVA, Tukey’s post hoc test and two-way ANOVA were used for statistical analysis (GraphPad PRISM v5.01). p<0.05 was considered to indicate statistical significance.

## Results

### Kv1.3 is palmitoylated at the immunological synapse in human CD4^+^ T lymphocytes

Following TCR recognition of peptide–MHC complexes, several key signaling proteins must localize to the membrane and lipid rafts at the immunological synapse. Reversible palmitoylation directs proteins such as Lck, Fyn, and LAT to these sites, where they play crucial roles in T-cell activation [16,17]. Similarly, the voltage-gated potassium channel Kv1.3 regulates the extent of lymphocyte activation, with hyperactive T cells displaying enhanced Kv1.3 currents. Pharmacological inhibition of Kv1.3 effectively restrains excessive T-cell activation, underscoring its role as a key modulator of immune responses [25–27]. Previous studies have demonstrated that Kv1.3 targets lipid rafts via association with caveolin [8]. However, most T cells lack caveolins, and the mechanisms underlying the localization of Kv1.3 in these microdomains remain uncertain [14]. In this context, palmitoylation, which is involved in a number of cellular events, has emerged as a key mechanism for protein recruitment into lipid rafts [28,29]. In fact, palmitoylation mediates the targeting of the regulatory subunit Kvb2.1 to lipid rafts and the IS in T cells during the immune response [15]. Therefore, we investigated whether Kv1.3 underwent palmitoylation in T lymphocytes. Using the Acyl Biotin Exchange (ABE) assay on human CD4^+^ T cells, we found that, similar to other Kv channels [30,31], Kv1.3 is palmitoylated (Fig. 1A). Furthermore, upon short incubation with anti-CD3/CD28 beads, T-cell activation increased the level of palmitoylation (Fig. 1A). In addition, the combination of a click-reaction with a proximity ligation assay (PLA) (Fig. 1B) demonstrated that S-acylated Kv1.3 localized at the plasma membrane of CD4^+^ T cells (Fig. 1C). Further analysis of human Raji-CD4^+^ T-cell conjugates demonstrated that the palmitoylated channel preferentially situated at the IS (Fig. 1D). To analyze the spatial distribution of the channel, CD4^+^ T cells were electroporated with Kv1.3-YFP mRNA and exposed to supported-lipid bilayers (SLBs) containing histidine tagged ICAM1, CD58, CD80, and UCHT1 Fab [4]. SMAC compartments were defined as indicated (Fig. 1E). Lymphocytes were fixed after a 15 min incubation at 37 °C with the SLB, click-reaction and PLA assays were performed, and the palmitoylated Kv1.3 was visualized by total internal reflection microscopy (TIRFM). The PLA signal (palmitoylated Kv1.3) displayed an annular distribution surrounding the cSMAC, suggesting that palmitoylation rearranged Kv1.3 into specific regions within the IS (Fig. 1F). Analysis of the sagittal XZ and YZ planes revealed that both nonpalmitoylated and palmitoylated channels exhibited IS staining patterns compatible with Kv1.3 internalization in the cSMAC (Fig. 1G).

**Fig. 1.**
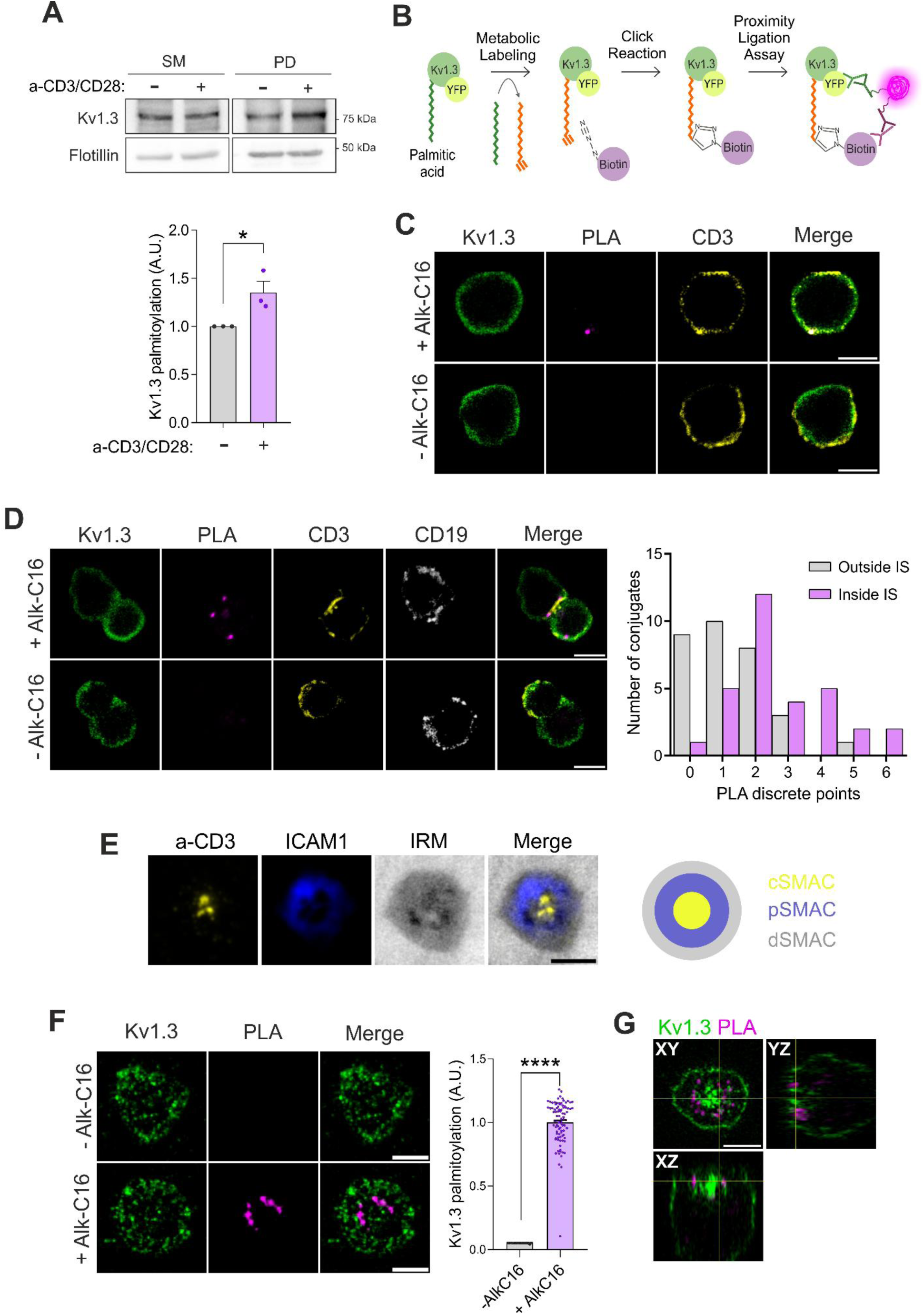
Kv1.3 is palmitoylated at the immunological synapse. (A) Palmitoylation assay (ABE) in CD4^+^ T cells in the absence (-) or presence (+) of anti-CD3/CD28 dynabeads for 15 min. Flotillin was used as a positive control. SM: starting materials, PD: pull-down. Bottom panel, quantification of Kv1.3 palmitoylation, which was calculated as the PD/SM ratio, and the control was normalized to 1 for better visualization. The data are presented as the mean ± SE of 3 independent blood donors. *p < 0.05 by Student’s t test. (B) Schematic of the click-chemistry and proximity ligation assay (PLA) technique. First, endogenous palmitic acid (green) was replaced by alkynyl palmitic acid (orange) through metabolic labeling. Next, biotin-azide was attached via a click reaction. Finally, specific antibodies against YFP (or Kv1.3 for the endogenous channel) and biotin conjugated with complementary probes were incubated with the samples for the PLA reaction. Therefore, the PLA signal indicates palmitoylated Kv1.3-YFP. (C) Representative confocal images of the PLA signal in human CD4^+^ T cells in the presence (+Alk-C16) or absence (-Alk-C16) of clickable palmitic acid. Following the PLA reaction, specific antibodies were used to label endogenous Kv1.3 (green) and CD3 for the plasma membrane (yellow). The PLA signal (magenta) shows palmitoylated Kv1.3. The merged panels show triple colocalization in white. Scale bars represent 5 μm. (D) Kv1.3 is palmitoylated at the immunological synapse (IS) between Kv1.3 YFP-electroporated CD4^+^ T cells and Raji B cells. Representative confocal images of the PLA in the presence (+Alk-C16) or absence (-Alk-C16) of palmitic acid. Following the PLA reaction, specific antibodies against Kv1.3 were used to identify endogenous channels (green), CD3 for T lymphocytes (yellow), and CD19 for B cells (white). The PLA signal (magenta) shows palmitoylated Kv1.3. The merged panels show triple colocalization in white. Scale bars represent 5 μm. Right panel, quantification of the PLA signal measured inside and outside the IS. (E) Representative TIRF images showing the marker distribution in synapses formed by CD4^+^ T cells on the SLB and a cartoon of the SMAC compartments. Anti-CD3 (a-CD3) in yellow accumulates at the central SMAC (cSMAC), and ICAM1 in blue forms a ring at the peripheral SMAC (pSMAC). Interference reflection microscopy (IRM) image showing the cell-SLB contact and delimits the distal SMAC (cSMAC) in gray. (F) Representative TIRF images of CD4^+^ T cells electroporated with Kv1.3 YFP forming synapses on the SLB. PLA assays revealed palmitoylated channels forming a ring in the presence (+Alk-C16) but not in the absence (-Alk-C16) of clickable palmitic acid. Scale bars represent 5 μm. Right plot, quantification of Kv1.3 palmitoylation on the basis of the PLA signal intensity. The data were normalized to the average of the +AlkC16 condition. The plot shows the mean ± SE of n > 50 cells from 3 independent blood donors. ****p < 0.0001 by Student’s t test. (G) Orthogonal views of a CD4^+^ T-cell forming an IS with an SLB and imaged by 3D Airyscan microscopy. Magenta, palmitoylated Kv1.3 (PLA signal); green, total Kv1.3. Note the internalization of the palmitoylated channel in the YZ and XZ planes.

Acyltransferases catalyze protein palmitoylation and have emerged as promising therapeutic targets in autoimmune disorders [32]. Altered Kv1.3 function has also been linked to autoimmune diseases [26]. Therefore, it is tempting to speculate that impaired channel palmitoylation could be associated with pathogenicity. Therefore, the identification of the acyltransferase that catalyzes Kv1.3 palmitoylation is crucial. Evidence suggests that the Ca^2+^-dependent acyltransferase ZDHHC21 palmitoylates the TCR in lymphocytes [33], as well as several proteins located in lipid rafts, such as the α1A-adrenergic receptor [34]. In this scenario, imaging of ZDHHC21 in synapse conjugates revealed that this enzyme is significantly recruited to the IS, paralleling the spatial distribution of Kv1.3 and suggesting a functional role in these regions (Fig. 2A–B). Interestingly, further analysis demonstrated that ZDHHC21 colocalized with both CD3 (TCR) and Kv1.3, particularly within the IS region (Fig. 2C–D). Finally, we knocked down ZDHHC21 in CD4^+^ cells and analyzed Kv1.3 palmitoylation. The results indicated that ZDHHC21 inhibition caused deficient Kv1.3 palmitoylation (Fig. 2 E–G).

**Fig. 2.**
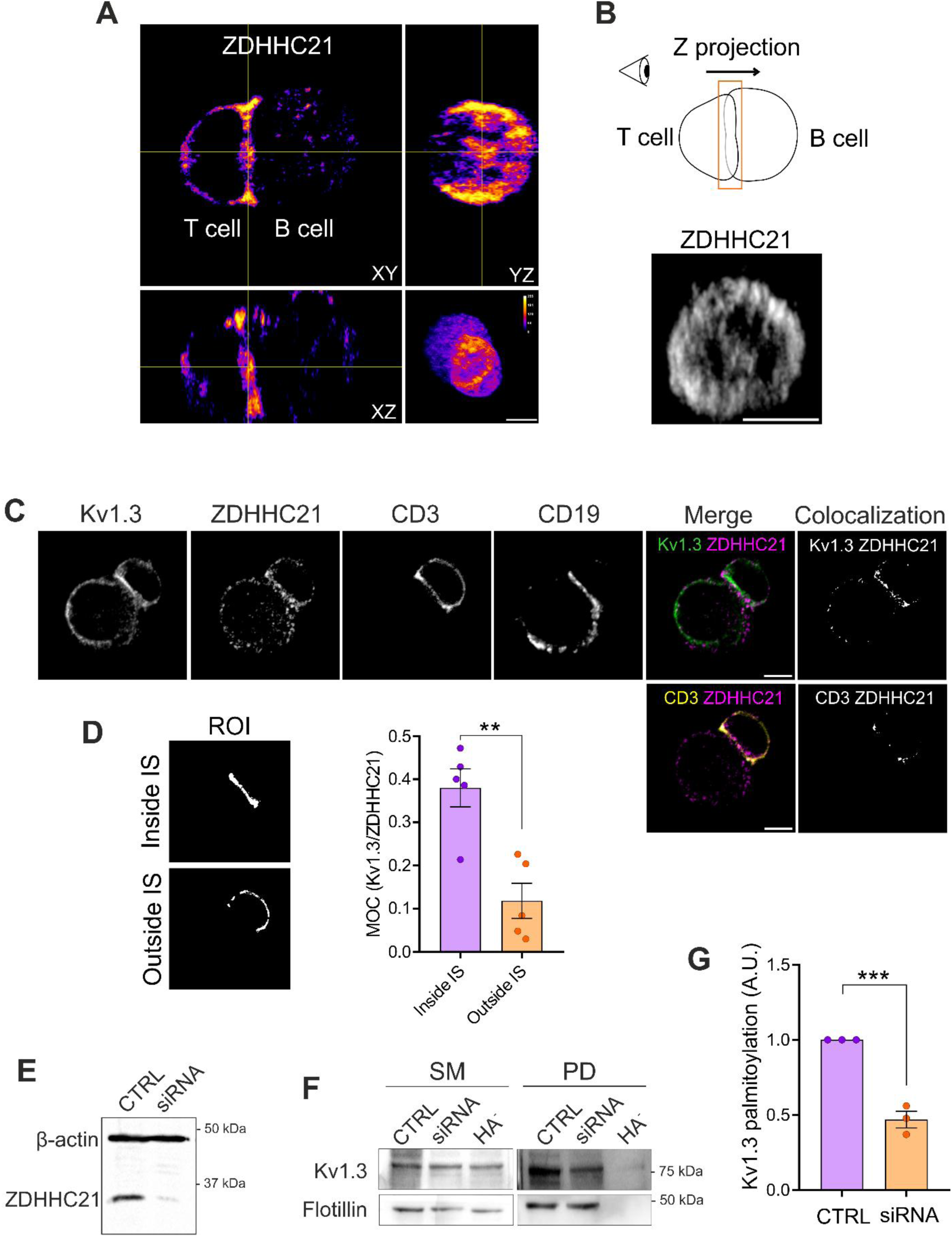
ZDHHC21 acyltransferase mediates Kv1.3 palmitoylation. (A) Orthogonal views of a representative synapse conjugate between a CD4^+^ T cell and a Raji B cell immunostained with an anti-ZDHHC21 antibody. Three-dimensional reconstruction revealed the distribution of the acyltransferases in both lymphocytes. (B) Top panel, schematic of the image processing for the visualization of ZDHHC21 at the synaptic platform in (A). In the bottom panel, intensities from the stacks corresponding to 3 µm of the synaptic region were compiled in a single Z projection. Note that, similar to Kv1.3, ZDHHC21 organizes as a distal ring with partial accumulation at the center of the synapse. The scale bar represents 5 μm. (C) Representative confocal images of synapse conjugates between CD4^+^ T cells and Raji B cells. Immunostaining of endogenous Kv1.3, ZDHHC21, CD3 (T cells), and CD19 (B cells) was performed. Merged panels show colocalization between Kv1.3 and ZDHHC21 (top) and between CD3 and ZDHHC21 (bottom). Colocalized pixels identified by Otsu’s automatic thresholding are displayed to facilitate visualization. The scale bar represents 5 μm. (D) Representative regions of interest (ROIs) inside and outside the IS regions were manually identified using the CD3 and CD19 masks. Right panel, Manders overlap coefficient (MOC) between Kv1.3 and ZDHHC21 inside and outside the IS. Data are presented as the mean ± SE of 5 cells from the same donor. **p < 0.01 by Student’s t test. (E) Representative Western blot of ZDHHC21-silenced CD4^+^ T cells. Lymphocytes were electroporated with ZDHHC21 siRNA, and after a 48-h incubation, whole-cell lysates were analyzed by Western blotting. β-actin was used as a loading control. CTRL: control, untreated cells. (F) Representative Western blot of Kv1.3 palmitoylation in CTRL- or ZDHHC21-knockdown (siRNA) CD4^+^ T cells. Flotillin was used as a positive control. SM: starting materials, PD: pull-down, HA^-^: negative control. (G) Quantification of Kv1.3 palmitoylation under basal conditions (CTRL) and after ZDHHC21 silencing (siRNA). The data were normalized to those of the CTRL. The data are presented as the means ± SEs of 3 independent donors. ***p < 0.001 by Student’s t test.

### Palmitoylation affects the spatial distribution and dynamics of Kv1.3 at the IS

Once the acyltransferase of Kv1.3 was identified, we investigated the functional consequences of palmitoylation on channel activity and its impact on lymphocyte physiology. ZDHHC21 knockdown was not a suitable model because this enzyme is responsible for the palmitoylation of a significant number of proteins, such as the TCR [33], potentially masking Kv1.3-specific effects. To overcome this limitation, we generated a Kv1.3 Cys*_less_* mutant in which all the intracellular cysteines were substituted with serine residues (Fig. 3A). We confirmed that Cys_less_ targeted normally to the plasma membrane in HEK293 cells (Fig. S1A–D). As expected, compared with the WT channel, the Cys*_less_* mutant showed resulted in significantly reduced palmitoylation levels in HEK293 cells (Fig. 3B–C). Similarly, the palmitoylation of Cys*_less_* was negligible in CD4^+^ T cells (Fig. 3D–F). Interestingly, TIRF imaging revealed notable colocalization of the PLA signal of WT Kv1.3 within the ICAM1 ring (Fig. 3E). Analysis of intermediate mutants—lacking N-terminal Cys (Nt_C-_ Kv1.3), lacking C-terminal Cys (Ct_C-_ Kv1.3), or with CSS-Palm v2.0 software-predicted Cys mutation (CSS Kv1.3)—revealed that palmitoylation of Kv1.3 occurs at multiple alternative and complementary cysteine residues distributed along the cytoplasmic loops and tail of the channel (Fig. S2A–B). Among these, palmitoylation at C-terminal cysteines may play a more prominent role in regulating channel abundance at the IS, whereas palmitoylation at the cysteines in the N-terminal region and the second intracellular loop may be more important for controlling subsynaptic distribution and modulating Kv1.3 function during T-cell activation. This finding was supported by the greater accumulation of the Ct_C-_ mutant at the IS (Fig. S2C) and the increased localization of Nt_C-_ and CSS Kv1.3 in CD58-enriched regions (Fig. S2D), accompanied by lower levels of the CD69 activation marker in the CSS mutant (Fig. S2E).

**Fig. 3.**
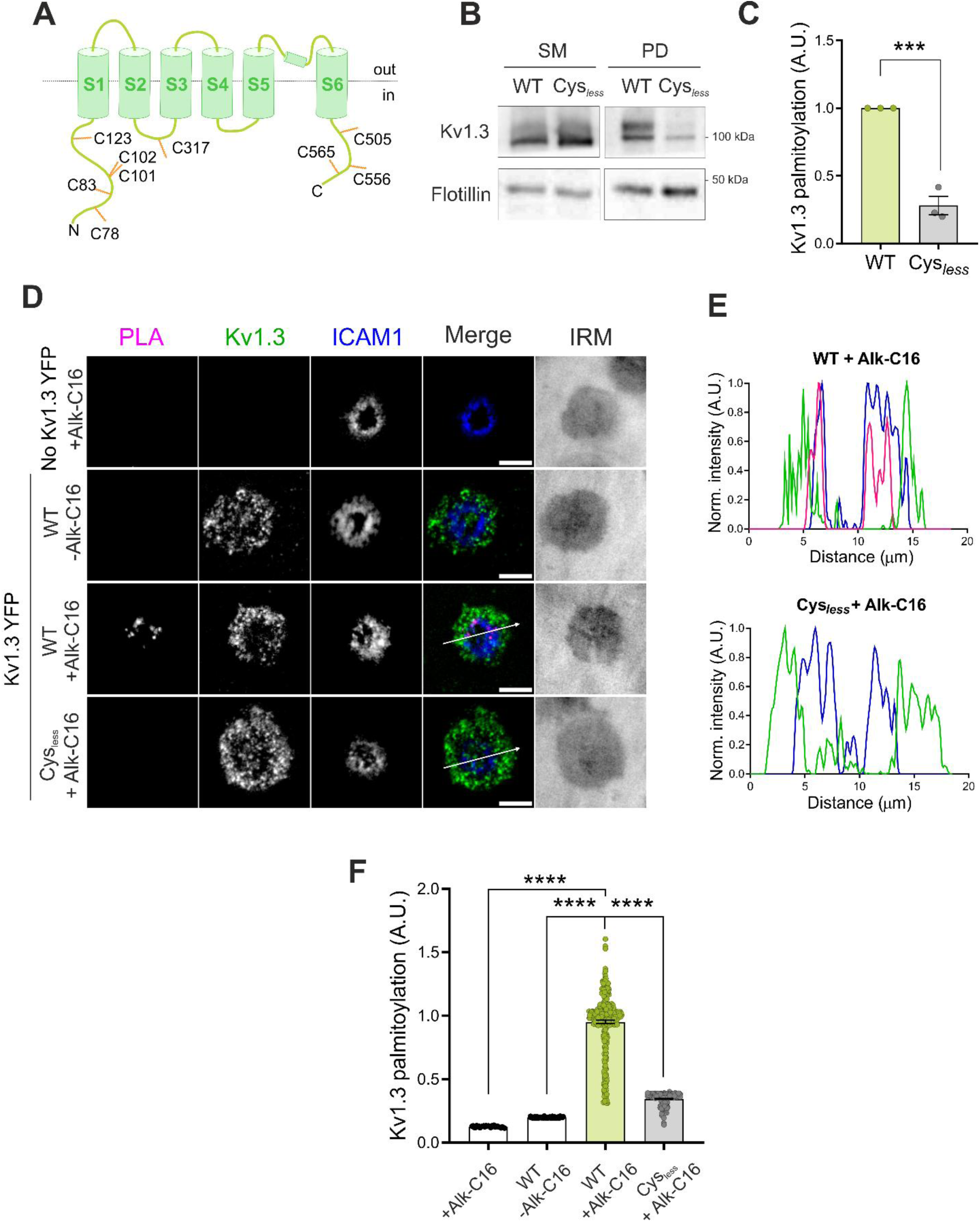
Characterization of the Kv1.3 Cys*_less_* palmitoylation mutant. (A) Schematic of Kv1.3 highlighting all the intracellular cysteines. (B) Representative Western blot of Kv1.3 palmitoylation in HEK293 cells transfected with wild-type (WT) or palmitoylation mutant (Cys*_less_*) Kv1.3 YFP. Flotillin was used as a positive control. (C) Quantification of the palmitoylation of the Kv1.3 channels. The palmitoylated fractions were relativized to the starting materials, and the data were normalized to those of the WT condition. The data are presented as the means ± SEs of 3 independent experiments. **p < 0.01 by Student’s t test. (D) Representative TIRF images of CD4^+^ T cells forming synapses on the SLB. T cells were either not transfected (no Kv1.3 YFP) or electroporated with WT or Cys*_less_* Kv1.3 YFP. T cells were incubated in the absence (-Alk-C16) or presence (+Alk-C16) of clickable palmitic acid. The PLA signal indicates the palmitoylated channel. The ICAM1 ring indicated the formation of a synapse. The merged panels show colocalization, and the IRM image shows the interference reflection microscopy image. Magenta, PLA; green, Kv1.3; blue, ICAM1. Scale bars represent 5 μm. (E) Pixel-by-pixel analysis of the PLA (magenta), Kv1.3 (green), and ICAM1 (blue) signals highlighted by arrows from the images in (D). (F) Quantification of the PLA signal intensity, which corresponds to Kv1.3 palmitoylation. Data were normalized to the average of the WT+Alk-C16 condition. Data are presented as the means ± SEs of n>50 cells from 3 different blood donors. ****p < 0.0001 according to one-way ANOVA with a post hoc Tukey test.

Next, we analyzed whether the ability of Cys*_less_* to target lipid rafts was altered, as palmitoylation regulates protein affinity for these membrane domains. Kv1.3 was localized within lipid raft microdomains in both Jurkat (Fig. 4A) and primary CD4^+^ T cells (Fig. 4B), as evidenced by the presence of channels in flotillin-enriched low-buoyant sucrose-density fractions and its colocalization with cholera toxin B (CTxB), a lipid raft marker, respectively. In this scenario, Kv1.3 was recruited to lipid rafts concentrated at the central regions of the IS in primary CD4^+^ T cells (Fig. 4C). In contrast, the Cys*_less_* mutant exhibited lower raft microdomain localization. Single-particle tracking analysis revealed that compared with the WT, Cys*_less_* channels exhibited higher speeds, displacements, and confinement ratios, which is consistent with a preferential distribution in less densely packed membrane domains. Notably, when trajectories were segregated by spatial location, the differences were slightly more pronounced in central tracks, where raft microdomains are typically enriched (Fig. 4D).

**Fig. 4.**
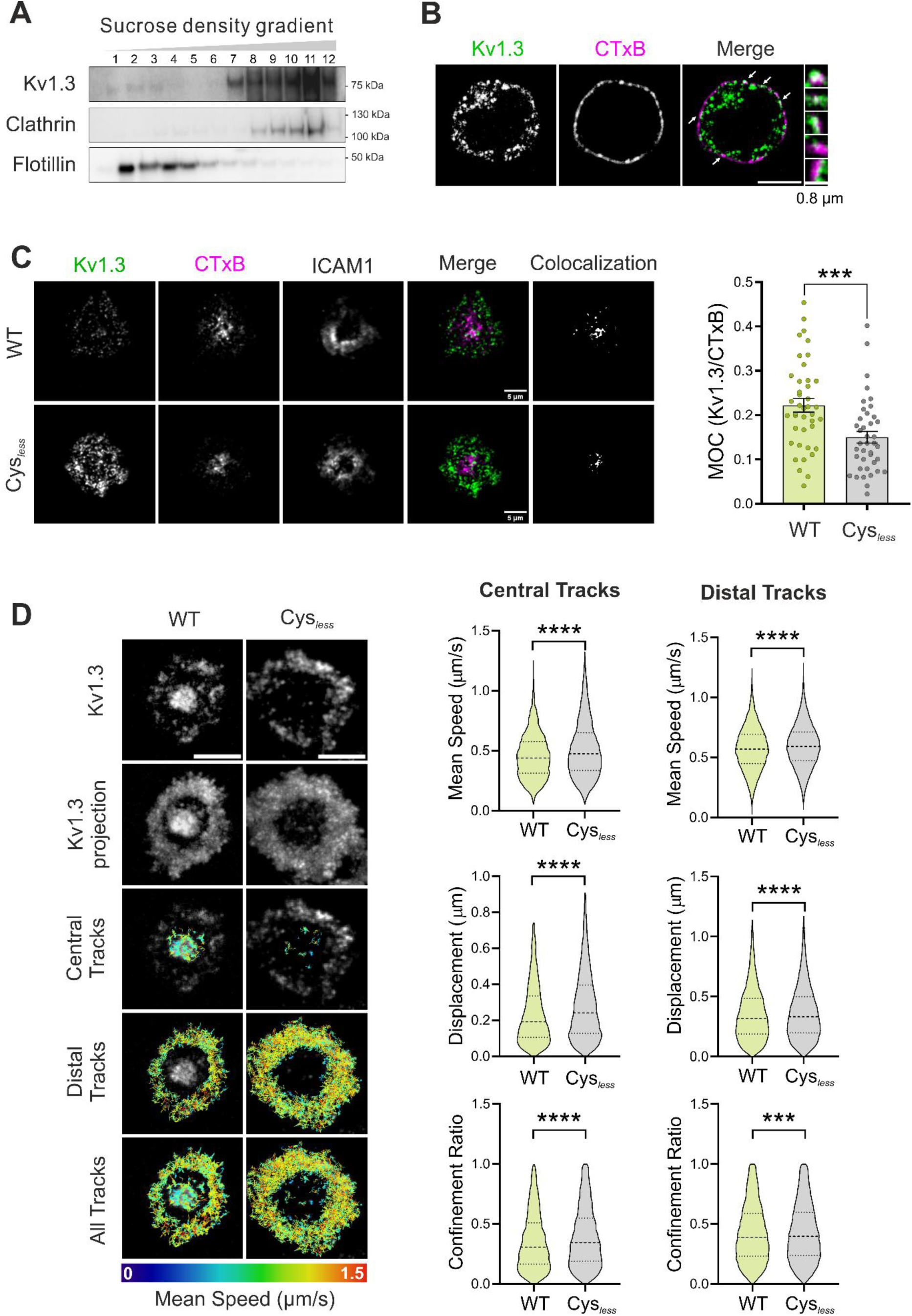
A lack of palmitoylation impairs the lipid raft accumulation of Kv1.3 at immunological synapses and increases channel membrane mobility. (A) Kv1.3 partially localizes to detergent-resistant fractions in Jurkat T lymphocytes. Cell lysates were separated by a sucrose gradient, and low-buoyant fractions from lower (1) to higher (12) densities were analyzed by Western blotting. Clathrin and flotillin were used as nonraft and raft markers, respectively. (B) Confocal images of a CD4^+^ T-cell showing colocalization between endogenous Kv1.3 (green) and lipid rafts (cholera toxin subunit B (CTxB), magenta). The merged panel shows colocalization in white. The arrows indicate the zoomed-in insets on the right. The scale bar represents 5 μm. (C) Representative TIRF images of CD4^+^ T cells electroporated with WT or Cys*_less_* Kv1.3 YFP (green) to form synapses on the SLB. Lipid rafts were stained with the B subunit of cholera toxin (CTxB, magenta). The ICAM1 ring indicated the formation of a synapse. Merge shows the Kv1.3 and CTxB channels. Colocalization panels show colocalization pixels identified by Otsu’s automatic thresholding for better visualization. Scale bars represent 5 μm. Right panel, Manders overlapping coefficient (MOC) of Kv1.3 inside CTxB regions. Data are presented as the means ± SEs of n > 40 cells from 3 independent blood donors. ***p < 0.001 by Student’s t test. (D) Live CD4^+^ T cells transfected with wild-type (WT) or Cys*_less_* Kv1.3 YFP were incubated with SLB at 37 °C and imaged for 1 min at 0.3 s/frame using TIRFM. Single-frame images of Kv1.3 and a time projection of all the frames recorded are shown. Single-particle tracking was performed. The track color indicates the mean speed of the track. The color scale bar on the bottom indicates the minimum values in blue and the maximum values in red in μm/s. Scale bars represent 5 μm. Right violin plots depict the quantification of the mean track speed (average velocity across all the links of the track, μm/s), total displacement (net distance traveled, μm), and confinement ratio (net displacement divided by total distance, 0 = fully confined, 1 = straight path). Data are the means ± SEs (n > 4000 trajectories) from 3 independent blood donors. ***p < 0.001 by Student’s t test.

### Palmitoylation participates in the centralization of Kv1.3 at the IS

To further explore the role of Kv1.3 palmitoylation in T cells, we studied the expression and spatial distribution of Kv1.3 Cys*_less_* at the IS. Human CD4^+^ T cells electroporated with either WT or Cys*_less_* Kv1.3 formed synapses on SLB. Unpalmitoylated Cys*_less_* channels increased in the IS (Fig. 5A–B). Next, we analyzed the specific distribution of Kv1.3 within each SMAC region. While the Cys*_less_* mutant preferentially localization to the dSMAC and pSMAC, its localization to the cSMAC was reduced by 50% compared with that of WT Kv1.3 (Fig. 5C–D). Notably, the cSMAC, a lipid raft-enriched region, is a site for the endocytosis of signaling complexes. Indeed, during T-cell activation, the TCR migrates from the dSMAC to the cSMAC, where it undergoes endocytosis to terminate signaling [35]. Similarly, we recently demonstrated that Kv1.3, initially located in the dSMAC, moves to the cSMAC for removal from the plasma membrane [4]. Evidence has demonstrated that Kv1.3 endocytosis, via a clathrin-dependent mechanism, requires channel ubiquitination [36–38]. In T cells, TCR ubiquitination has previously been associated with the cSMAC and can lead to either internalization or budding in extracellular vesicles, depending upon the clathrin adapter recruited [39,40]. Therefore, we studied the ubiquitination of Kv1.3 in synapse conjugates using the PLA technique. The endogenous Kv1.3 channel was ubiquitinated at the central region of the synaptic interface, which is compatible with the cSMAC (Fig. 5E). To further analyze the ubiquitination of Cys*_less_,* transfected HEK293 cells were incubated with Phorbol 12-myristate 13-acetate (PMA) to induce PKC-dependent ubiquitin-mediated internalization of the channel [36,38]. Our results revealed that the Cys*_less_* mutant, which was less recruited to the cSMAC, underwent less ubiquitination under endocytic triggers, which was consistent with a longer persistence at the IS dSMAC (Fig. 5F–G). Thus, palmitoylation regulates the migration of Kv1.3 to the cSMAC, the primary site of ubiquitination and removal from the plasma membrane.

**Fig. 5.**
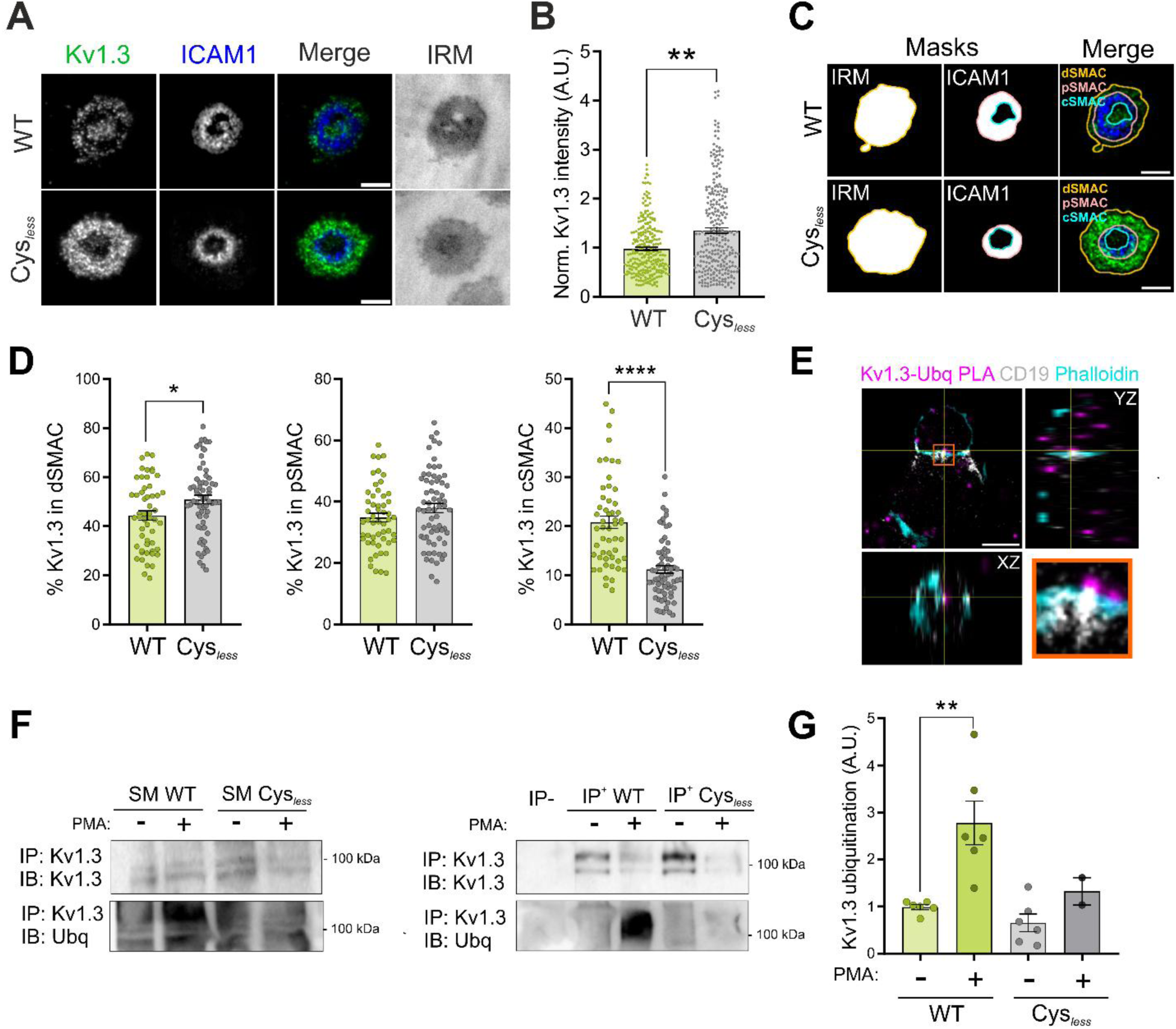
Palmitoylation facilitates the ubiquitination and central accumulation of the channel at the immunological synapse. (A) Representative TIRF images of CD4^+^ T cells electroporated with WT or Cys*_less_* Kv1.3 YFP to form synapses on the SLB. The ICAM1 ring indicates the formation of a synapse. The merged image shows Kv1.3 (green) and ICAM1 (blue) expression. IRM, interference reflection microscopy. Scale bars represent 5 μm. (B) Quantification of Kv1.3 intensity at the immunological synapse normalized to the average of the WT for each donor. Data are presented as the means ± SEs of n > 300 cells from 3 independent blood donors. **p < 0.01 by Student’s t test. (C) ROIs of each supramolecular activation complex (SMAC) in (A). IRM images were used to define the total cell contact area. The ICAM1 ring indicates the pSMAC. IRM regions outside or inside the ICAM1 ring were classified as dSMAC or cSMAC, respectively. (D) Kv1.3 intensity was quantified within each ROI corresponding to the cSMAC, pSMAC, dSMAC, and the entire synapse. The percentage of Kv1.3 intensity in each SMAC was calculated with respect to the total number of synapses. Data are presented as the means ± SEs of n > 50 cells from 3 independent blood donors. *p<0.05, ****p < 0.0001 by Student’s t test. (E) A PLA was performed using anti-ubiquitin and anti-Kv1.3 antibodies to detect ubiquitinated Kv1.3. Orthogonal views from a representative confocal image of ubiquitinated Kv1.3 (PLA signal, magenta) in a synapse conjugate between a human CD4^+^ T cell and a Raji B cell. CD19 (gray) was used as a B-cell-specific marker, and phalloidin (cyan) was used to stain the actin filaments. The PLA signal detected within the synaptic contact is located at the center of the synapse (XZ plane). (F) Ubiquitination assay in HEK293 cells transfected with either WT or Cys*_less_* Kv1.3 YFP. Cells were incubated in the absence (-) or presence (+) of phorbol 12-myristate 13-acetate (PMA) for 30 min to induce protein internalization as a positive control. Cell lysates were immunoprecipitated (IP) for Kv1.3 and immunoblotted (IB) for both Kv1.3 and ubiquitin (Ubq). SM: starting materials, IP: immunoprecipitation, IP^-^: negative control in the absence of antibody. (G) Quantification of channel ubiquitination. The data are presented as the means ± SEs of at least 2 independent experiments. **p < 0.01 by Student’s t test.

Kv1.3 internalization is a ubiquitin-dependent mechanism involving clathrin-mediated endocytosis and trafficking through the endosomal pathway [36–38]. Therefore, we next analyzed Kv1.3 localization at early endosomes and the plasma membrane (PM) in HEK293 cells. As expected, upon PMA-induced endocytosis, the colocalization of WT Kv1.3 with early endosomes increased (Fig. 6A–B). Conversely, the Cys*_less_* mutant exhibited impaired endocytosis, as evidenced by reduced colocalization with EEA1 and an elevated abundance at the cell surface (Fig. 6A–B). Concomitantly, in contrast to that in the WT, the amount of Kv1.3 Cys*_less_* present at the plasma membrane, as shown by the membrane protein biotinylation assay, remained stable in the presence of PMA (Fig. 6C–D).

**Fig. 6.**
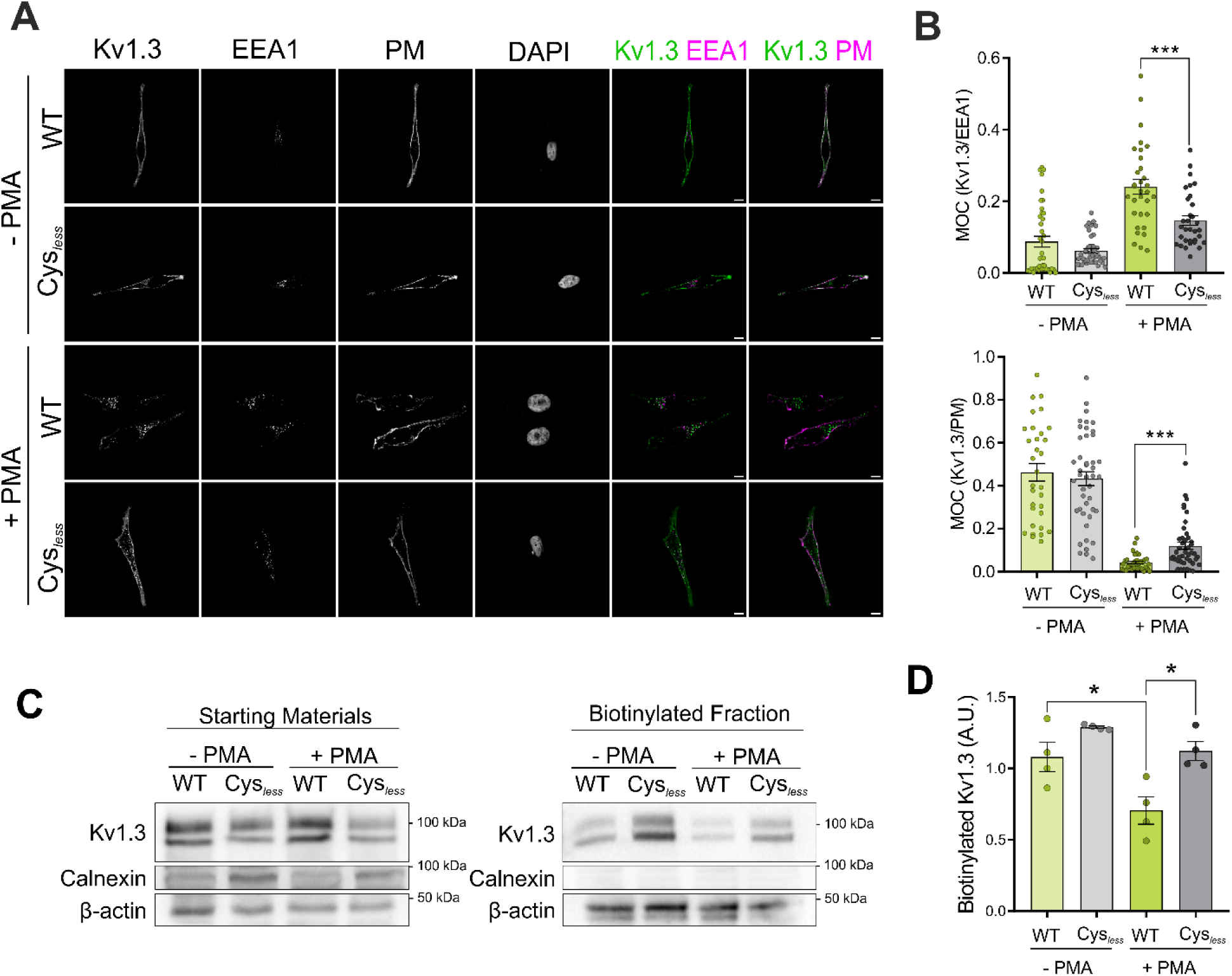
Palmitoylation facilitates Kv1.3 endocytosis. (A) Representative confocal images of HEK293 cells transfected with either WT or Cys*_less_* Kv1.3 YFP. The cells were incubated in the absence (-PMA) or presence (+PMA) of PMA for 30 min, after which early endosomes (EEA1) were identified. The plasma membrane (PM) and nuclei were stained with wheat germ agglutinin (WGA) and DAPI, respectively. The merged panels show the indicated colocalizations in white. Green, Kv1.3 channels; magenta, cell markers (EEA1 or PM). Scale bars represent 10 μm. (B) Manders overlapping coefficients between Kv1.3 and EEA1 (top panel) and between Kv1.3 and the PM (bottom panel). Data are presented as the means ± SEs of n > 30 cells from 3 independent experiments. ***p < 0.001 by one-way ANOVA with post hoc Tukey test. (C) Membrane protein biotinylation assay of HEK293 cells transfected with either WT or Cys*_less_* Kv1.3 YFP and incubated in the absence (-PMA) or presence (+PMA) of PMA for 30 min to trigger endocytosis. Calnexin was used as a negative control because it is a transmembrane protein located in the ER and is absent at the PM, and β-actin was used as a loading control. (D) Quantification of Kv1.3 biotinylation in arbitrary units (A.U.). The data are presented as the means ± SEs of 4 independent experiments. *p < 0.05 by one-way ANOVA with post hoc Tukey test.

We have demonstrated that the absence of palmitoylation affects the spatial distribution of Kv1.3, impairing channel endocytosis. The internalization of Kv1.3 is essential for terminating the immune response. Excessive Kv1.3-related signaling leads to persistent activation and proinflammatory responses [41]. Thus, endocytosis fine-tunes the precise balance of functional surface channels. In this sense, the C-terminal domain of Kv1.3 contains a number of interacting clusters, which are involved in channel trafficking and membrane stabilization (Fig. 7A). Thus, scaffold proteins, such as PSD95 and cortactin, stabilize the channel at the cell surface, preventing endocytosis [36,42]. Therefore, we evaluated whether the level of palmitoylation altered the channel association with PSD95 and cortactin. WT and Cys*_less_* Kv1.3 colocalized with CTxB and PSD95 in HEK293 cells. PMA triggered massive endocytosis of WT Kv1.3, which was partially prevented in the Cys*_less_* mutant. While PMA disrupted the colocalization of PSD95 with WT Kv1.3, significant colocalization of PSD95 with Cys*_less_* remained (Fig. 7B). The persistent colocalization of Kv1.3 Cys*_less_* and PSD95 was further supported by major coimmunoprecipitation of both proteins (Fig. 7C). While the PDZ domain (PSD95 binding domain) is located at the most distal part of the Kv1.3 C-terminus, the cortactin binding site is situated between two main forward trafficking motifs (Fig. 7A) [42,12]. Evidence has demonstrated that the cortactin binding domain is vital for channel immobilization at the plasma membrane of T cells. Cortactin anchors the channel to the actin cytoskeleton, promoting channel stability at the immunological synapse [42]. In this context, we analyzed whether palmitoylation could also affect cortactin binding. TIRF images of CD4^+^ T cells that formed an IS on the SLB revealed that a significant portion of the channels were located within the actin ring (Fig. 7D). Furthermore, coimmunoprecipitation assays demonstrated that, similar to PDS95, the Cys*_less_* mutant exhibited greater cortactin association than the WT channel did in HEK293 cells (Fig. 7E). Therefore, impaired palmitoylation increased the association of channels with membrane-stabilizing adaptor proteins, such as PSD95 and cortactin, preventing centralization, ubiquitination and removal from the plasma membrane.

**Fig. 7.**
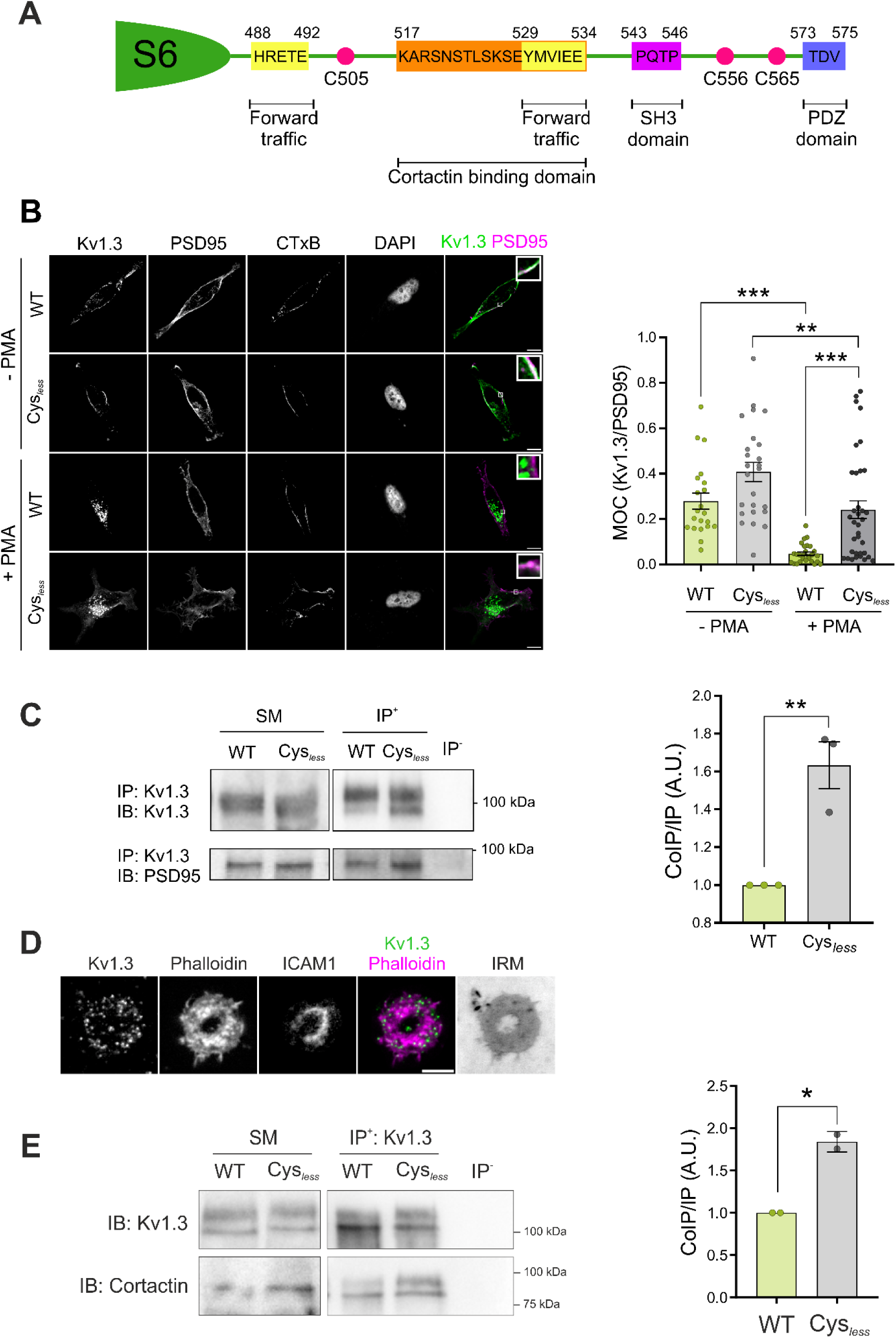
The absence of palmitoylation altered channel‒protein interactions with plasma membrane-stabilizing proteins in HEK293 cells. (A) Illustration of the C-terminal domain of human Kv1.3 highlighting several known scaffold protein binding domains, cysteine residues, and anterograde traffic signatures. (B) Representative confocal images of HEK293 cells transfected with either WT or Cys*_less_* Kv1.3 YFP. Cells were incubated in the absence (-PMA) or presence (+PMA) of PMA for 30 min to induce endocytosis. Cells were stained for PSD95 and the B subunit of cholera toxin (CTxB, a lipid raft marker). Nuclei were stained with DAPI. The merged panels show colocalization in white, as magnified in the enlarged inset. Green, Kv1.3; magenta, PSD95. Scale bars represent 10 μm. The right panel shows the Manders overlap coefficient between Kv1.3 and PSD95. Data are presented as the mean ± SE of n>20 cells from 3 independent experiments. **p < 0.01, ***p < 0.001 by one-way ANOVA with a post hoc Tukey test. (C) Left panel, representative immunoblot of the coimmunoprecipitation between Kv1.3 and PSD95. HEK293 cells were transfected with either WT or Cys*_less_* Kv1.3 YFP. Cell lysates were immunoprecipitated (IP) against Kv1.3 and immunoblotted (IB) against Kv1.3 and PSD95. SM, starting materials; IP^+^, positive immunoprecipitation; IP^-^, negative control in the absence of antibody. Right panel, quantification of the results of the coimmunoprecipitation of Kv1.3-PSD95 by normalizing the coimmunoprecipitation of PSD95 (CoIP) to the immunoprecipitation (IP) of Kv1.3. The data are presented as the means ± SEs of 3 independent experiments. **p < 0.01 by Student’s t test. (D) Representative TIRF images of human CD4^+^ T cells electroporated with Kv1.3 YFP forming a synapse on an SLB. Phalloidin labeled the actin ring. The merged panel shows colocalization. Green, Kv1.3; magenta, phalloidin. The scale bar represents 5 μm. IRM shows interference reflection microscopy images. (E) Coimmunoprecipitation (CoIP) of Kv1.3 with cortactin. HEK293 cells were transfected with either wild-type (WT) or Cys*_less_* Kv1.3. Cell lysates were immunoprecipitated (IP) against Kv1.3 YFP and immunoblotted (IB) against GFP (Kv1.3) and cortactin. SM: starting materials, IP^+^: positive immunoprecipitation, IP^-^: negative control in the absence of antibody. Right panel, quantification of the coimmunoprecipitation (CoIP) between Kv1.3 and cortactin by relativizing the CoIP of cortactin to the IP of Kv1.3. The data are presented as the means ± SEs of 2 independent experiments. *p < 0.05 by Student’s t test.

### Kv1.3 palmitoylation shapes channel activity, influencing T-cell physiology

The importance of protein palmitoylation in regulating the immune response has just started to emerge. Palmitoylation also affects crucial aspects of tumorigenesis and antitumor immunity [43]. Although the spatial organization of palmitoylated proteins during T-cell signaling is still poorly understood, alterations in the lipidation of this protein have been observed at the onset of immune-related diseases; therefore, palmitoylation may be at the forefront in the diagnosis and treatment of immune disorders [43,18]. Alterations in Kv1.3 palmitoylation affect protein dynamics, localization and channel–protein interactions. Therefore, we investigated the effects of palmitoylation on Kv1.3-related functional consequences during T-cell activation. Although the Cys*_less_* mutant was more abundant at the plasma membrane, its voltage-dependent K⁺ currents were reduced by nearly 50% (Fig. 8A), suggesting that palmitoylation regulates channel function through mechanisms beyond surface expression. The reduced function of the Cys*_less_* Kv1.3 correlated with an impairment in early T-cell activation. The activation of TCR signaling, as assessed by pZAP70 levels, was significantly reduced in CD4^+^ T cells expressing the Cys*_less_* mutant during synapse formation on the SLB. A time course analysis of pZAP70 staining revealed that T-cell activation was decreased from the initial stages of IS formation and remained low throughout the study (Fig. 8B–D). The physiological impact on early T-cell activation was further confirmed by Ca^2+^ imaging. Ca^2+^ flux is an important hallmark of T lymphocyte activation following IS formation. CD4^+^ T cells expressing endogenous Kv1.3, electroporated or not with either WT or Cys*_less_* channels, were loaded with the Calbryte™ Ca^2+^ indicator and monitored during the formation of ISs with Raji B-cell conjugates. The peak amplitude, upon TCR engagement, revealed that Ca^2+^ mobilization was significantly lower in the Cys*_less_* mutant (Fig. 8E, Online Resource 1).

**Fig. 8.**
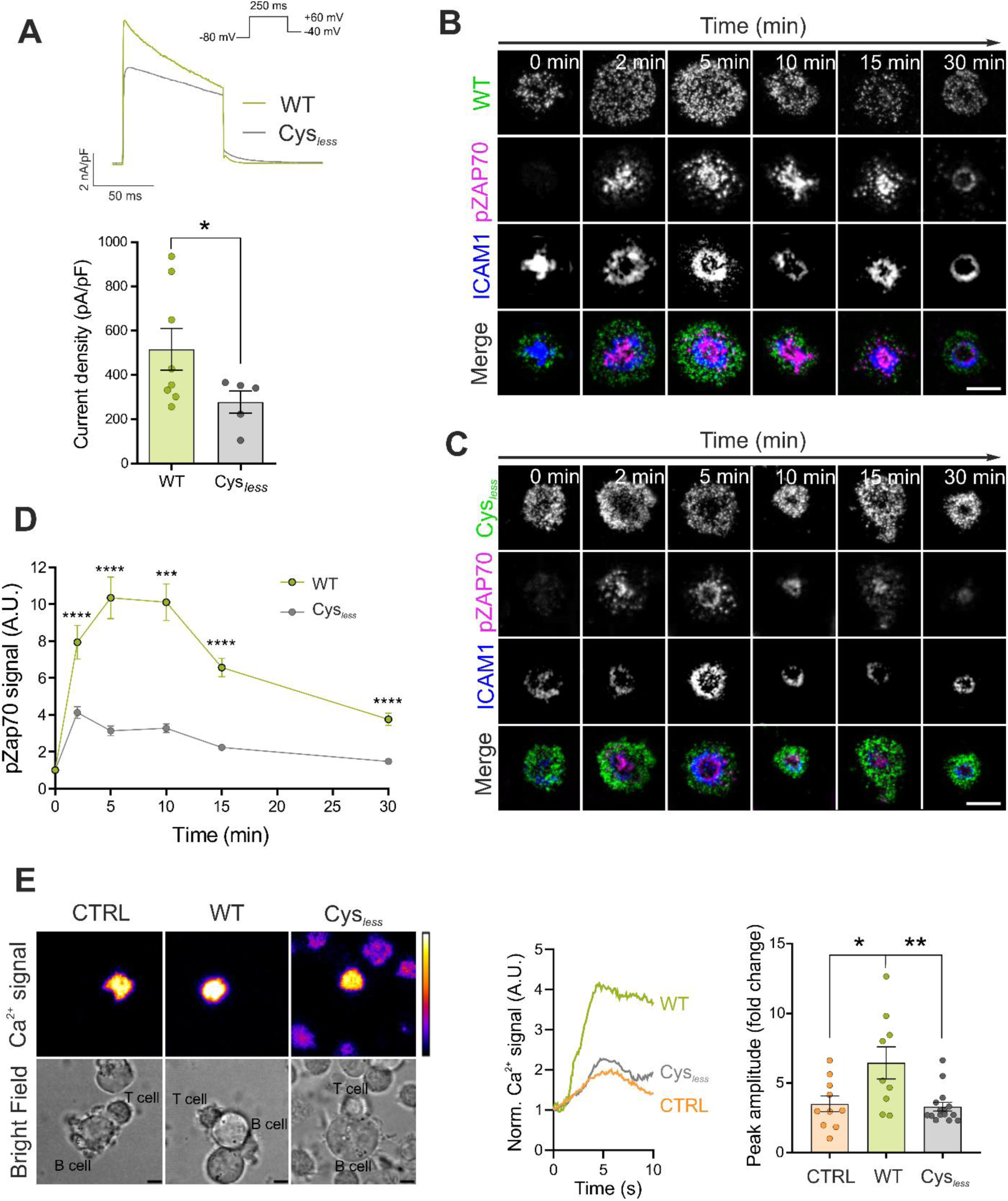
Functional consequences of the absence of Kv1.3 palmitoylation. (A) HEK293 cells were transfected with wild-type (WT) and Cys*_less_* mutant Kv1.3, and voltage-dependent currents were analyzed by patch clamp. Cells were held at -80 mV, and voltage-gated currents were elicited by 250 ms square pulses at +60 mV. Current densities were calculated (bottom panel). The data are presented as the means ± SEs of at least 5 independent cells per condition. *p < 0.05 by Student’s t test. (B–C) Representative TIRF images of CD4^+^ T cells electroporated with wild-type (WT) (B) or Cys*_less_* (C) Kv1.3 YFP and exposed to SLBs at different times (0, 2, 5, 10, 15, and 30 min). ICAM1 rings identify immunological synapses. pZAP70 staining indicated T-cell activation. The merged panels show colocalization. Green, Kv1.3 channels; magenta, pZAP70; blue, ICAM1. Scale bars represent 5 μm. (D) Quantification of pZAP70 intensity at different times of IS formation. The data were normalized to the average of the 0 min time point for each donor. Data are presented as the means ± SEs of n > 100 cells. ***p < 0.001, ****p < 0.0001 by Student’s t test. (E) Left panel, representative snapshots of Ca^2+^ signaling recordings of CD4^+^ T cells forming immunological synapses with Raji B cells. T lymphocytes expressing endogenous Kv1.3 (CTRL) or electroporated with either wild-type (WT) or Cys*_less_* Kv1.3 YFP were loaded with Calbryte™ 630 AM and incubated with Raji B cells. The calibration bar shows the signal intensity, and the scale bars represent 5 μm. Center panel, representative Ca^2+^ traces of the 3 conditions. Right panel, quantification of the maximum peak amplitude calculated as the maximum Ca^2+^ intensity relative to the initial signal. Data are presented as the means ± SEs of n > 9 cells. *p < 0.05, **p < 0.01 by Student’s t test.

## Discussion

Kv1.3 regulates T-cell activation and proliferation by sustaining calcium flux at the immunological synapse. In this study, we characterized how distinct PTMs, palmitoylation and ubiquitination integrate to regulate Kv1.3 function and localization within the IS, thus modulating its role in early T-cell activation. Additionally, we identified ZDHHC21 as the acyltransferase responsible for Kv1.3 palmitoylation, presenting a novel potential therapeutic target for immune disorders. Upon T-cell activation, Kv1.3, initially localized within the dSMAC, migrates to the lipid raft-enriched cSMAC where it is removed from the plasma membrane to terminate the activating signal. Previous studies have reported that ubiquitination is essential for Kv1.3 clathrin-mediated endocytosis [37,36]. Our results suggested that S-acylated Kv1.3 underwent ubiquitination, increasing its recruitment to cSMAC and facilitating channel removal from the plasma membrane. In this sense, Kv1.3 palmitoylation plays a crucial role in the termination of Kv1.3 function, likely by regulating its association with scaffold proteins, such as PSD95 and cortactin, interactions that reduce ubiquitination and removal from the plasma membrane. Moreover, S-acylated cysteines throughout the Kv1.3 sequence have compensatory effects. Overall, palmitoylation emerges as a critical cellular mechanism for regulating the biology of the Kv1.3 channel and remodeling T-cell function during the initiation of the immune response. These findings highlight the potential of targeting palmitoylation—either at the channel level or the entire machinery—as a promising therapeutic method for developing pharmacological strategies targeting immunological disorders.

Kv1.3 is aberrantly expressed in immune cells during autoimmune and lymphoblastic maladies [25,26]. Channel blockade ameliorates the prognosis of these diseases in animal models [44–47], and channel inhibitors have proven effective for the treatment of certain autoimmune disorders in humans [48]. Kv1.3 accumulates at the IS [7], and its compartmentalization at the synapse is altered during autoimmune processes [27]. However, the underlying mechanisms remain obscure. We introduce Kv1.3 palmitoylation as a key regulator of channel dynamics and membrane arrangement at the IS. Palmitoylation allows highly dynamic regulation of protein affinity for membrane microdomains [49]. This posttranslational modification is an essential regulator of many signaling pathways, including those involved in TCR-induced T-cell activation. In fact, palmitoylation constitutes a recognized mechanism for lipid raft targeting, in which proteins are in close proximity to interact and modulate signaling pathways [29]. Hence, the palmitoylation of CD4 and CD8 coreceptors regulates raft affinity and enhances coreceptor activity [50,28]. Similarly, the palmitoylation of LAT is required for lipid raft localization and the mediation of T-cell activation by this protein [51]. Kv1.3 is an S-acylated raft-associated protein in T cells. The localization of Kv1.3 in membrane rafts modulates channel gating properties in T cells [52]. Palmitoylation regulates several ion channel proteins [53], including the *Shaker* channels Kv1.1 [54] and Kv1.5 [55] and the regulatory Kvb2.1 subunit [15]. Here, we demonstrate that palmitoylation serves as a complementary mechanism for targeting Kv1.3 to lipid rafts in caveolin-deficient T cells and modulating channel currents.

Palmitoylation is catalyzed by zinc finger DHHC domain-containing (ZDHHC) acyl transferases. The human ZDHHC family comprises 23 members that have been linked to various diseases. Hence, these enzymes have attracted much interest as promising candidates for drug development in recent years [32]. In this work, we identified ZDHHC21 as the acyl transferase responsible for the palmitoylation of the channel, revealing a novel potential therapeutic target. This enzyme, which also S-acylates the TCR and TRPV2 channels [33], paralleled the distribution of Kv1.3 at the IS. Moreover, ZDHHC21 was recruited to the IS, where it colocalized with Kv1.3 and CD3, suggesting local activity. Consequently, Kv1.3 palmitoylation at the IS determines the spatial distribution of the channel. Kv1.3 is uniformly distributed in the plasma membrane of human CD4^+^ T cells. Upon TCR engagement, Kv1.3 is radially excluded from the center of the IS, localizing at the dSMAC. However, a small but significant portion of Kv1.3 accumulated locally at the cSMAC. The channel is endocytosed via a clathrin-dependent mechanism in this region to attenuate Kv1.3 activity during the IS [4]. Protein kinase Cθ (PKCθ) is recruited at the cSMAC [56]. Kv1.3 is a target for PKC phosphorylation, which induces channel ubiquitination and clathrin-dependent endocytosis [36,38]. Thus, it is tempting to speculate that Kv1.3 targets the cSMAC undergoing PKCθ and clathrin-dependent endocytosis, which represents a mechanism for controlling Kv1.3 activity, buffering Ca^2+^ signaling and/or termination of Kv1.3 activity. In fact, our PMA results, as a model of PKC activation, demonstrated that the palmitoylation-deficient Kv1.3 Cys*_less_* mutant underwent less endocytosis through a mechanism that involves low ubiquitination and strong interactions with PSD95 and cortactin. Therefore, we propose that the level of Kv1.3 palmitoylation alters its association with scaffold proteins. This is not a minor issue because MAGUK proteins, such as PSD95, are located at the IS and stabilize proteins at this spot, serving as hubs for protein—protein interactions [57,13,58]. In the case of Kv1.3, PSD95 binds to the distal region of the C-terminus, in close proximity to the cortactin-binding domain [42,13]. Moreover, cortactin participates in the actin-mediated immobilization of Kv1.3 [42]. In this context, we hypothesized that the enhanced binding of Cys*_less_* Kv1.3 to cortactin would increase channel immobilization at the dSMAC by the anchorage to the actin ring, thus preventing channel internalization. The increased association of Kv1.3 Cys*less* with cortactin validated this hypothesis. Hence, by enduring its association, PSD95 and cortactin fine-tune the spatial distribution and internalization of the channel. The lasting persistence of Kv1.3 would trigger an exacerbated channel-related signal, resulting in significant immune outcomes that lead to prolonged responses and inflammatory disorders.

In conclusion, our work provides the first evidence of the palmitoylation-dependent effects of Kv1.3 on T-cell physiology. We demonstrated that Kv1.3 is palmitoylated in lymphocytes and identified ZDHHC21 as the responsible acyltransferase. Moreover, our results suggest a highly dynamic interplay between Kv1.3 palmitoylation and channel localization within different SMACs at the IS. Impaired Kv1.3 palmitoylation leads to increased channel accumulation at the IS dSMAC through decreased ubiquitination and removal from the plasma membrane in the cSMAC. Mapping the dynamic regulation of Kv1.3 offers new perspectives beyond genetic studies, providing critical insights into its involvement in a wide repertoire of pathophysiological processes.

## Acknowledgments

We acknowledge the CCiTUB, the Kennedy Trust for Rheumatology Research, the Institute for Developmental and Regenerative Medicine, and Carl Zeiss GMBH for the microscopy facilities used in this research. We would also like to thank all the anonymized blood donors who contributed to our study.

## Author contributions

Conceptualization: MNP, MPV, MLD, AF, JC

Methodology: MNP, MPV, ABG, JC

Investigation: MNP, MPV, ABG, JC

Visualization: MNP, MPV, JC

Supervision: MLD, AF, JC

Writing—original draft: MNP, AF, JC

Writing—review & editing: MNP, MPV, ABG, MLD, AF, JC

## Data and materials availability

All data are available in the main text or the supplementary materials.

