## Supplemental Data for "Kv1.3 palmitoylation regulates spatial distribution and channel removal from the immunological synapse"

**Supplementary Information**

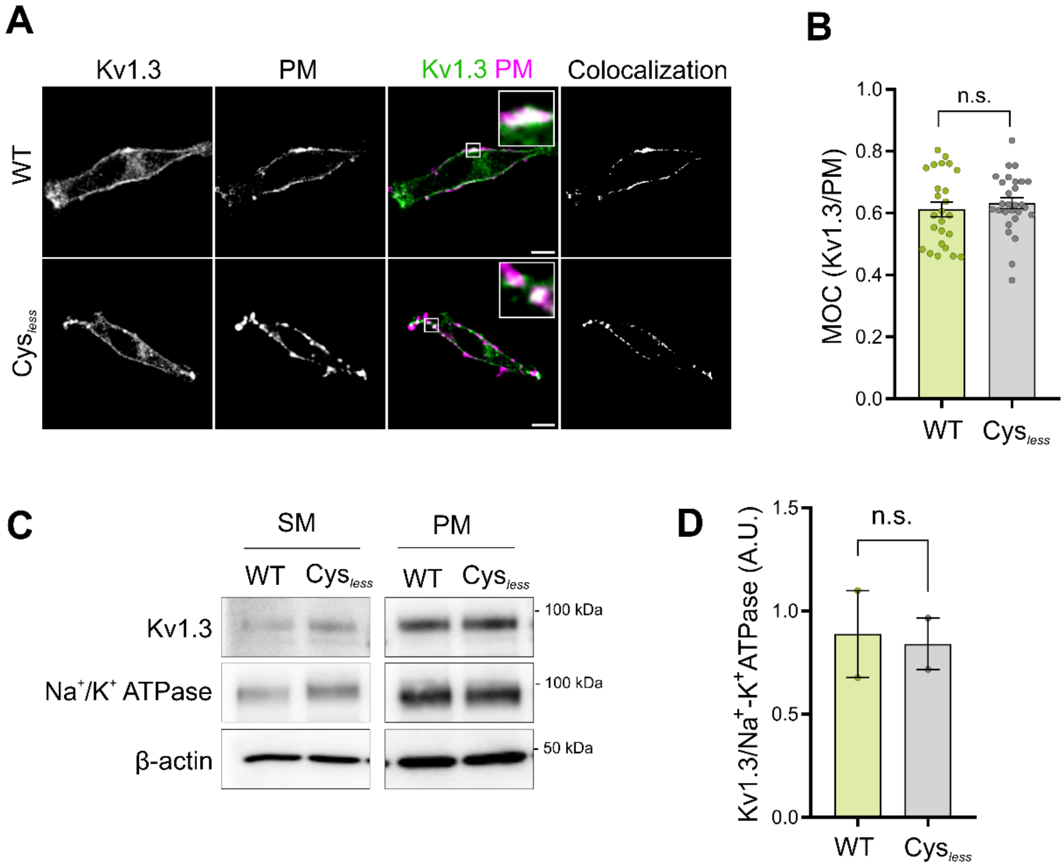


**Fig. S1** The Cys*less* Kv1.3 channel preserves normal membrane targeting. HEK293 cells were transfected with either wild-type (WT) or Cys*less* Kv1.3 YFP, and cell surface expression was analyzed. (A) Representative confocal images of Kv1.3 targeting the cell surface. The plasma membrane (PM) was stained with WGA. The merged panels show colocalization in white. Green, Kv1.3 channels; magenta, PM marker. Scale bars represent 10 µm. (B) Manders overlapping coefficient (MOC) of Kv1.3 and the PM. The data are presented as the means ± SEs of 3 independent experiments. Student's t test indicated no significant (n.s.) differences. (C) The plasma membranes were isolated, and the expression of Kv1.3 was analyzed. $Na^+/K^+$ ATPase was used as a plasma membrane marker, and β-actin was used as a loading control. SM, starting material; PM, plasma membrane fraction. (D) The expression of Kv1.3 at the plasma membrane was normalized to the $Na^+/K^+$ ATPase intensity. The data are means ± SEs of 2 independent experiments. Student's t test indicated no significant (n.s.) differences.

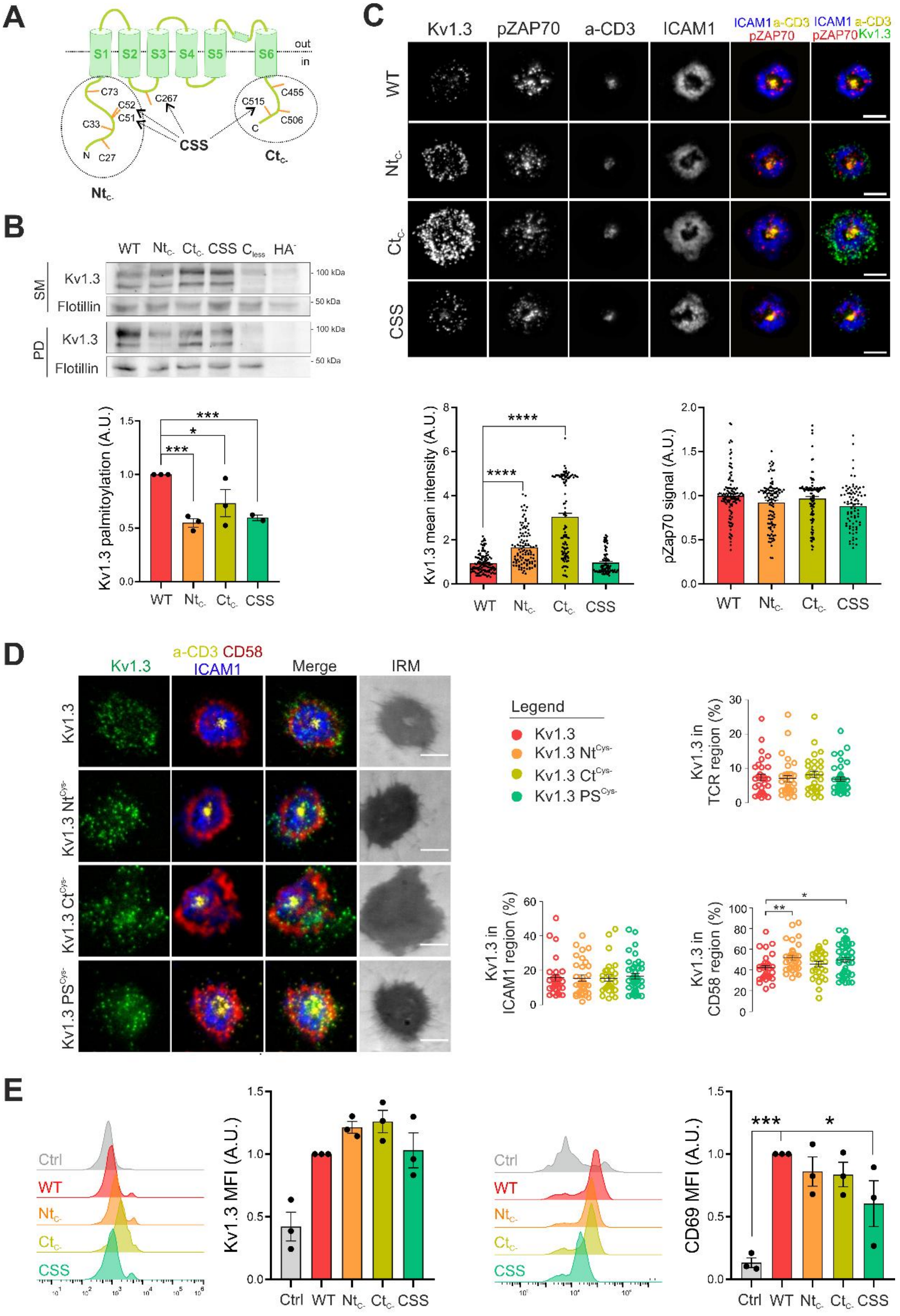
A
out
in
S1
S2
S3
S4
S5
S6
C73
C52
C51
C33
C27
N
C267
CSS
C515
C455
C506
C
Ntc-
Ctc-
B
WT
Ntc-
Ctc-
CSS
Cless
HA-
SM
Kv1.3
Flotillin
PD
Kv1.3
Flotillin
100 kDa
50 kDa
Kv1.3 palmitoylation (A.U.)
***
*
***
WT
Ntc-
Ctc-
CSS
C
Kv1.3
pZAP70
a-CD3
ICAM1
ICAM1 a-CD3 pZAP70
ICAM1 a-CD3 pZAP70 Kv1.3
WT
Ntc-
Ctc-
CSS
Kv1.3 mean intensity (A.U.)
****
****
pZap70 signal (A.U.)
D
Kv1.3
a-CD3 CD58 ICAM1
Merge
IRM
Kv1.3
Kv1.3 NtCys-
Kv1.3 CtCys-
Kv1.3 PSCys-
Legend
Kv1.3
Kv1.3 NtCys-
Kv1.3 CtCys-
Kv1.3 PSCys-
Kv1.3 in TCR region (%)
Kv1.3 in ICAM1 region (%)
Kv1.3 in CD58 region (%)
**
*
E
Ctrl
WT
Ntc-
Ctc-
CSS
Kv1.3 MFI (A.U.)
CD69 MFI (A.U.)
***
*

**Fig. S2** Multiple cysteines are involved in Kv1.3 palmitoylation. (A) Cartoon of rKv1.3 highlighting all the intracellular cysteines. Specific Kv1.3 mutant channels were generated by substituting cysteine with serine at the intracellular N-terminal (circle, $Nt_{C-}$) and C-terminal (circle, $Ct_{C-}$) domains and those predicted by CSS-Palm 2.0 software (arrows, CSS). In addition, all the intracellular cysteines were mutated in the $Cys_{less}$ mutant. (B) Representative results of the palmitoylation acyl-biotin exchange assay. HEK293 cells were transfected with all Kv1.3 channels, and a palmitoylation assay was performed. Flotillin was used as a positive control. SM, starting material; PD, pull-down; $HA^-$, negative control. The quantification of Kv1.3 palmitoylation (bottom panel) was performed by calculating the PD/SM ratio and normalizing to the WT. The data are presented as the means ± SEs of 3 independent experiments. *$p < 0.05$, ***$p < 0.001$ by Student's t test. (C) Representative TIRF images of $CD4^+$ T cells electroporated with rKv1.3 YFP mutants forming synapses on SLBs for 15 min. CD3 and ICAM1 labeling indicated the formation of an IS. pZAP70 measured early TCR signaling. The merged panels show the colocalization of the different markers. Green, Kv1.3; red, pZAP70; yellow, CD3; blue, ICAM1. Scale bars represent 5 µm. The bottom panels show the quantification of the recruitment of Kv1.3 channels to the IS (left) and the pZAP70 signal intensity (right). The intensity value of each cell was normalized to the mean value of the WT condition. Data are presented as the means ± SEs of $n > 60$ cells from 2 independent blood donors. ****$p < 0.0001$ according to one-way ANOVA with a post hoc Tukey test. (D) Representative TIRF images of human $CD4^+$ T cells incubated with SLB and fixed 15 min post-incubation. Cells were transfected with WT rKv1.3 YFP or its mutants (green). The merged panel shows the overlap of the green, red, yellow and blue channels. The IRM shows interference reflection microscopy images. Scale bars represent 5 µm. The color code of each mutant is shown on the right. The percentages of Kv1.3 abundance in the a-CD3 (UCHT1), CD58, and ICAM1 regions were calculated (right panels). Data are presented as the means ± SEs ($n > 30$). *$p < 0.05$, **$p < 0.01$ according to one-way ANOVA with post hoc Tukey test. (E) Flow cytometry results showing the Kv1.3 (left panels) and CD69 (right panels) intensities. The mean fluorescence intensities (MFIs) were normalized to that of the WT (1) for better visualization. The data are presented as the means ± SEs of 3 independent blood donors. *$p < 0.05$, ***$p<0.001$ according to Student's t test.
